# Redundancy relocated: the distributed metaparalog pool of an Arctic *Pelagibacter*ales population

**DOI:** 10.64898/2026.10.07.757358

**Authors:** Ana-Belen Martin-Cuadrado, Riccardo Rosselli, Francisco Rodriguez-Valera

## Abstract

The presence of metaparalogs (related genes diverging in function) has been proven to be extensive in the pangenomes of streamlined Pelagibacterales genomes. Here by leveraging a large number of complete genomes from an Arctic population we have been able to establish their universality and mechanistic origins. The diversity of some metaparalog families was as high within a single population than comparing with another (Mediterranean) population/species, which points to an ancient diversification of metaparalog families predating genomospecies separation. Metaparalog families provided diversity for transport and envelope diversity which would provide vast capacities to degrade substrates within DOM and interact with other cells or viruses or other subcellular entities (like EVs). The pool does not saturate at fifteen genomes. Our results highlight the immense importance of the pangenome in understanding the biology of bacteria in natural environments and how critical is the study of population genomics for the future of microbiology.

**Highlights:**

- A “common goods” genomic strategy for coexisting cells has been found in a streamlined bacterium.
- The pangenome is distributed among cells so that together can efficiently degrade marine dissolved organic matter.
- Synteny and distribution reflect ancient adaptation.
- The pangenome is an asset for the population as a whole.

**eTOC blurb:** Martin-Cuadrado et al. show that 15 Arctic SAR11 isolates from one water sample share most of their flexible genes as divergent homologs (metaparalogs) held at conserved chromosomal slots, so that functional redundancy lost from each streamlined genome is hugely amplified by the population.

## INTRODUCTION

The SAR11 clade constitutes up to 50% of marine bacterioplankton in global surface waters, representing perhaps the most abundant bacterial group in Earth’s oceans.^1^ The evolutionary success of SAR11 has been attributed partially to genome streamlining, with small genome sizes (1.2-1.7 Mb), minimal non-coding DNA, and elimination of seemingly redundant genes.^2^ This genomic minimalism was thought to reflect optimization for nutrient-limited marine environments where small genomes and cells provide a better surface-to-volume ratio, a critical parameter for osmotrophic heterotrophs. One symptom of streamlining is the scarcity of paralogs. Gene paralogs are multiple copies of homologous genes within genomes and typically provide functional diversification.^3^ Their existence has been taken as a model for genomic innovation in eukaryotes, where gene duplication is extremely frequent.^4^ However, their prevalence in prokaryotic genomes came as a surprise.^5^ In *Escherichia coli* K12, ca. 28% of the genes have a homolog elsewhere in the genome sharing at least 30% amino acid identity. Defined by this threshold, paralogs are much less prevalent in *Pelagibacter* or other streamlined aquatic microbes^6^ (see Supplementary Information).

However, recent pangenomic analyses have revealed a paradox. In the Pelagibacterales genomospecies Ia.3.VII (also known as gMED),^7^ which is very prevalent in near-surface Mediterranean offshore waters,^8^ populations maintain substantially flexible genomes with extensive within-population redundancy (genes clustered at ≥30% amino acid identity). These redundant genes, distributed among the coexisting cells of a population, have been termed “metaparalogs”.^9,10^ Metaparalogs act as a proxy for having a large paralog collection, as large prokaryotic genomes do,^11^ albeit distributed across different genomes within the population (a local pangenome). Each strain-specific region in gMED was associated with a specific gene pool whose members, though divergent, retained mechanistically related, even if biologically distinct, function.^12^ Thus, the divergent transporters of the TRAP and TTT families could in several cases be assigned different putative substrates on the basis of the catabolic genes flanking them.^9^ Likewise, the three ammonium transporters of the gMED reference genome, sharing only 29% aminoacid similarity, differ not in fold but in domain architecture, one carries a fused phosphoprotein phosphatase domain, which points to a sensory rather than a transport role. Within each transporter family, multiple sequence variants with equivalent substrate-binding domains but differing specificities constitute metaparalog sets; for example, the nine TRAP (DctPQM) and the nine TTT (TctCBA) variants identified in gMED.^9^ Metaparalogs have been interpreted as multiplying the metabolic network of a population while leaving individual genomes and cells small enough to perform their role in the highly diluted environment of the open ocean (or similar oligotrophic environments) in which a surface/volume ratio is a critical feature. This perspective depicts the pangenome as a “common good” on which the ecological efficiency of the population depends.^10^

However, the approach followed by Molina-Pardines et al. using single amplified genomes (SAGs) and long-read metagenomes to define the limits of the flexible genes (flexome), could not relate a metaparalog to the cell that carries it. A long read (average size 7.4 kb)^9^ establishes that a gene is present in the population, in its immediate genomic context. SAGs are often incomplete and the relative location of contigs in them hard to establish. Consequently, the divergence between two metaparalogs has never been measurable against the divergence between the genomes carrying them.

Recently, a set of 16 Arctic pelagibacters belonging to genomospecies Ia.1 have been isolated and sequenced.^13,14^ They were all isolated from a single sample of water under the sea ice, allowing us to replicate a similar analysis to that carried out by Molina-Pardines in the Mediterranean (a local pangenome) but with pure-culture complete genomes. Taking advantage of this dataset, we can compare metaparalog divergence within the context of the complete genome. In addition, we can approach the subject of synteny to establish where in the chromosomes metaparalogs reside, i.e., dispersed across the backbone, or concentrated in the spatially limited variable regions, as has been shown to recur across the Pelagibacteraceae genera.^15^ Lastly, synteny can also allow some inference relative to the mechanistic origin of the metaparalog pool and of the pangenome at large.

## RESULTS

### The pangenome of 15 Arctic *Pelagibacter* isolates

To study metaparalogs, it is critical to define first the core and flexible fractions in the pangenome of the strain set. From the original set of 16 isolates belonging to the Pelagibacterales subclade Ia.1 described by Sadler et al.,^13,14^ we selected 15 on the grounds of quality criteria (Tables S1 and S2; see Methods), and their pangenome (core and flexible) was re-analyzed. The Arctic genomes were uniformly streamlined, averaging 1.37 ± 0.05 Mb and 1,464 ± 53 genes (Table S1). Despite this uniformity in size and gene number, the population carries a substantially flexible genome. Clustering the 21,964 proteins at ≥80% amino acid identity and over ≥70% of the length of both sequences yielded 5,105 orthologous groups. Of these, 880 were present in at least 14 of the 15 genomes (691 of them in all fifteen) and were classified as core genome, whereas the remaining 4,225 constituted the flexible fraction (flexome). Under that framework, 13,013 of the 21,964 proteins (59.2%) belonged to core clusters (that is, had homologs at ≥ 80% identity in ≥14/15 strains) and 8,951 (40.8%) to the flexome (averaged over 15 genomes, this corresponds to 868 ± 10 core and 597 ± 50 flexible genes per strain). Core size varied predictably across prevalence and clustering-identity thresholds without altering the overall pangenome structure (Table S3).

These numbers differ from those reported by Sadler et al.^14^ for the 16 strains with an Anvi’o/MCL workflow, who reported 3,596 total gene clusters, 1,037 core clusters, and 1,661 singleton clusters. Our methodology returns similar values only when the identity is lowered to about 56%, and the coverage requirement is dropped. However, this criterion would not be suitable for the question addressed here. A bitscore-based threshold merges proteins that align over only part of their length, and it is precisely those partial-length relationships that a study of metaparalogy must resolve. The same caution applies when comparing to the Mediterranean *Pelagibacter* gMED genomospecies, analyzed by Molina-Pardines et al.,^9^ whose gene pool was delimited from a mixture of incomplete SAGs, and its diversity was recovered from long-read metagenomes.

### Most of the flexome in Arctic strains is made of metaparalogs

We define a metaparalog as a flexome gene with at least one homolog encoded in a different genome of the population, at ≥30% amino acid identity over ≥70% of the length of both sequences. A gene is assigned to the flexome when its 80% identity cluster is present in fewer than fourteen genomes, regardless of how similar its homologs may be; the two criteria are independent, and metaparalogs therefore span the full identity range from 30% to 100%. Of the 8,951 flexome genes, 7,424 (82.9%) met this criterion and fall into the 897 families analysed below. Per strain, metaparalogs made up 83.0 ± 5.7% of the flexome (range 71.7–91.4%). This is somewhat lower than the 95.7% reported for gMED,^9^ although the two figures are not comparable for the reasons given above. The remaining 1,527 genes are singletons with no detectable cross-genome relatives even at this permissive threshold.

We clustered metaparalogs into 897 families, whose distribution across strains is shown in Figure 1A. Family size ranged from two to more than 20 proteins, with families of 11 to 15 members being the most numerous (Figure 1B). Within families, identity falls monotonically with family size (Figure 1C), from a median of 93% in families of two proteins to 37% in those exceeding 20 members. Genes sharing more than 80% amino acid identity could be considered core of the corresponding set of strains, but absent from others, which makes them flexome and metaparalogs after our definitions. However, their high similarity makes functional diversification unlikely. The identity distributions become bimodal in the larger classes (16-20 and >20 members, Figure 1C). Families defined at 30% identity therefore combine two kinds of relationships, near-identical alleles of the same gene distributed across strains, and remote relatives close to the detection limit with likely divergent function. At the gross level of COG classification, transporters, metabolism (likely associated with transporters), and cell envelope categories dominate and increase with the bimodal nature of the family (Figure 1E).

**Figure 1.**
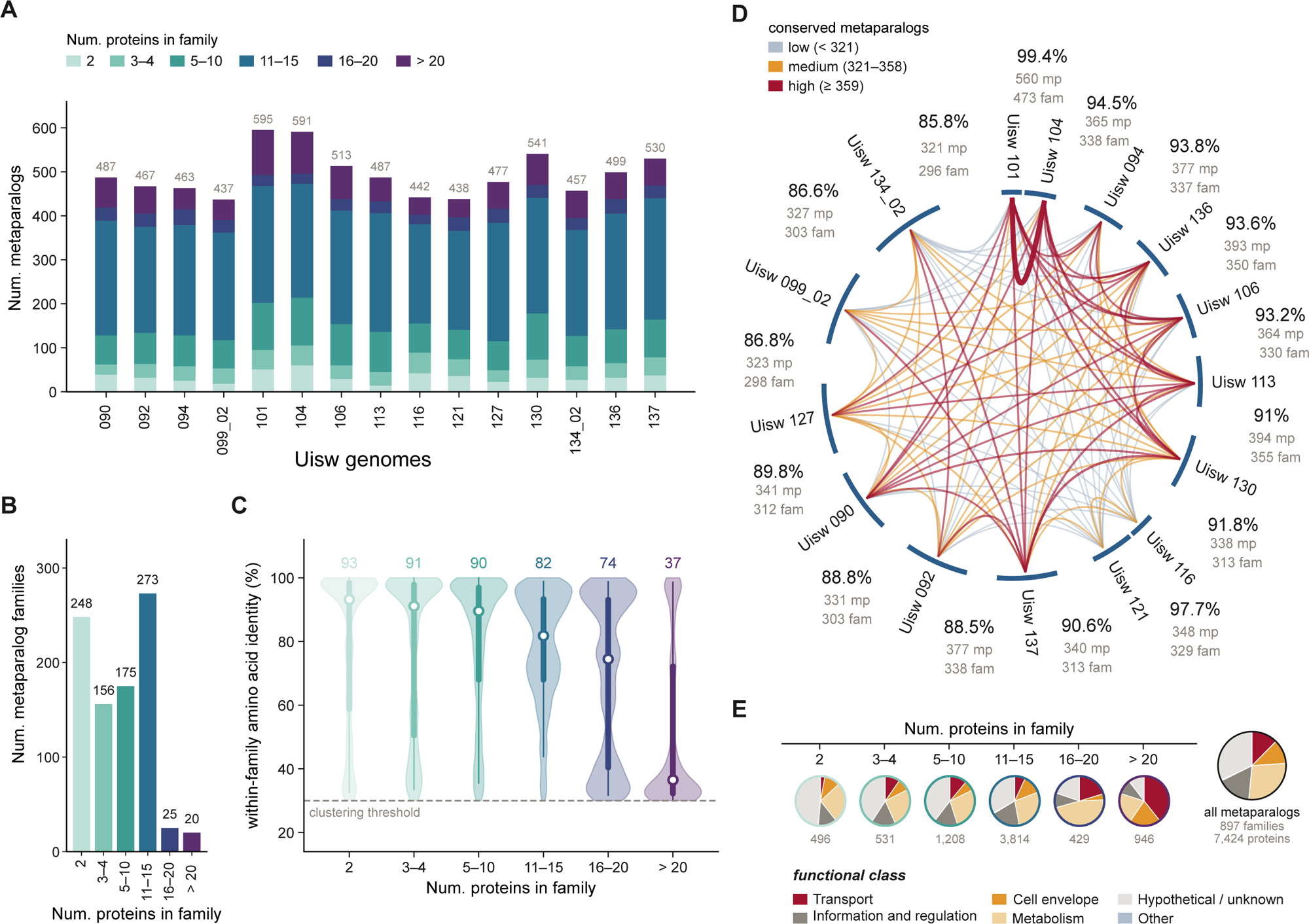
Metaparalog families of the Arctic *Pelagibacter* population. **(A)**, Number of metaparalog genes per genome, partitioned by the size of the family. (**B)**, Number of families in each size class. (**C)**, Pairwise amino acid identity between members of the same family, by family size. Violins show the full distribution; the point marks the median, the thick bar the interquartile range, and the thin bar the 5th to 95th percentiles. Median identity is given above each violin. (**D)**, Metaparalogs conserved between strains. Strains are arranged in order of ANI, and the angular gap between neighbours scales with their divergence. Values between each adjacent pair indicate ANI, conserved metaparalogs per genome (mp), and conserved metaparalog families (fam). Chords connect all strain pairs, coloured and scaled by the number of conserved metaparalogs. (**E)**, Functional composition of each size class (outline colour classified as in (C) and of all metaparalogs (right). The number of proteins is given beneath each pie. Proteins were assigned hierarchically to transport (transporter Pfam domain or annotation), cell envelope (COG category M), metabolism (COG C, E, F, G, H, I, P, Q), information and regulation (COG J, A, K, L, B, D, Y, V, T, U, O), hypothetical or unknown (no annotation, or COG S or R), and other.

Metaparalog families shared between pairs of strains ranged from 251 to 473 (median 312). Because the genomes are complete, it was possible to test whether this sharing is determined by genome-wide relatedness (Figure 1D). Relatedness explained only about a quarter of the variation (r = 0.522, r² = 0.27; r = 0.419 after controlling for flexome size), and the remaining were not explained by how closely the two strains are related. The two most closely related strains, Uisw_101 and Uisw_104, which share the same O-chain gene cluster (Figure S1) and an average nucleotide identity (ANI) 99.4% (they belong to the same clonal lineage), share by far the most of any pair, 473 families. The second most highly related pair, Uisw_116 and Uisw_121 (ANI 97.7%), share only 329 families, and each shares fewer families with the remaining thirteen genomes than any other strain (mean of 279 in both cases). On the other hand, the second-ranked pair (Uisw_130 and Uisw_137) share 365 families with an ANI of only 90.8%. Near-identity of the core genome is therefore compatible with either a large or an unremarkable shared pool, and a pair separated by 9.2% of genome-wide ANI divergence can share more families than one four times closer; i.e., the likelihood of sharing a metaparalog family is only weakly related to ANI except in the near-clonal pair Uisw_101 and Uisw_104. We next asked whether metaparalogs experience different selective constraints from the core genome. Pairwise *d*N/*d*S was estimated using the Nei–Gojobori and Yang–Nielsen methods for every pair of metaparalogs belonging to the same family and occupying the same syntenic position in different genomes, and compared with every pair of single-copy core orthologues. To minimize the effect of synonymous-site saturation, only comparisons with 0< *d*S < 1 were retained. Metaparalogs showed substantially weaker constraint than core genes but remained predominantly under purifying selection. Median *d*N/*d*S was 0.136 for metaparalog pairs (*n* = 20,574), twice the 0.064 observed for core genes (*n* = 65,162) (Figure S2A, Table S4). Thus, metaparalogs do not evolve free of constraint; they remain under purifying selection, although less strongly than the genes required by essentially every cell.

An independent analysis based on reciprocal best hits provided the same conclusion. Of 33,959 between-genome metaparalog pairs, 30,336 met stringent criteria for comparison (≥100 aligned codons and 0.01 ≤ *d*S ≤ 2.0), yielding a median *d*N/*d*S of 0.155. Of these, 95.1% had *d*N/*d*S < 0.5, whereas only six pairs (0.02%) exceeded one. A Yang–Nielsen sensitivity analysis of 18,651 stably estimated pairs gave a median of 0.084 and strongly agreed with the Nei–Gojobori estimates (Spearman’s ρ = 0.900; Figure S2B,C). The few gene-wide estimates exceeding unity therefore provide no robust evidence for pervasive positive selection that could be indicative of genes presently undergoing diversification.

### Metaparalog *loci* are not insulated from recombination

Recombination is extensive in natural SAR11 populations, including among strains recovered from individual seawater samples.^16^ We therefore asked whether the sequence variants occupying metaparalog *loci* are maintained by reduced genetic exchange. If so, metaparalog *loci* should show less evidence of recombination and greater concordance with the strain phylogeny than core genes.

PHI and NSS were jointly significant (P ≤ 0.05) in 45.9% of 690 single-copy core alignments and in 34.5% of 501 metaparalog *loci* (Fisher’s exact test, P = 8 × 10⁻⁵; Figure S2D; Table S4A,C). Part of this difference reflects statistical power, since *loci* present in fewer than ten genomes were detected least often (25.5%), but a modest deficit remained after accounting for the number of genomes and informative sites (odds ratio 0.61, P = 0.005). Gene trees, in contrast, gave no sign that metaparalog *loci* follow the strain phylogeny more closely than core genes. Mean gene concordance factors across the twelve internal branches of the reference phylogeny were 26.0% for 690 core-gene trees and 26.6% for 508 metaparalog-locus trees (Wilcoxon signed-rank P = 0.52; Figure S2E; Table S4D).

High concordance was restricted to the two most closely related genome pairs, reaching 91.7% for core trees and 95.4% for metaparalog trees for Uisw_101–Uisw_104 and 80.4% and 76.0% for Uisw_116–Uisw_121, whereas support for deeper branches ranged from only 2% to 30%. Thus, fewer than one-third of individual gene trees support 10 of the 12 branches of the concatenated core-genome topology. This genome-wide pattern extends the high recombination previously reported in SAR11 strains^16–18^ and indicates that metaparalog *loci* are not insulated from the genetic exchange affecting the rest of the genome.

### Metaparalogs occupy many small conserved positions

Because all fifteen assemblies are closed circular chromosomes, the distributed pool can be placed in its chromosomal context (Figure S3A). Partitioning each chromosome into maximal runs of consecutive flexible genes bounded by core genes yielded 3,055 variable regions, a mean of 204 per genome. Half of them (1,523, 49.9%) contain a single gene, the median region is 2 genes long, and only 69 exceed 15 genes, a proportion close to that reported for gMED (47.5%)^9^ and for coastal Hawaiian isolates.^15^ Metaparalogs are enriched in the smallest regions, accounting for 88.6% of the genes in regions of fewer than six genes against 75.5% in larger ones (Fisher exact test, odds ratio 2.53, *P* = 1.5 × 10⁻⁵⁹), and they are spread along the entire chromosome, whose densest tenth holds only 11.5–16.0% of them (Figure S3B).

Large genomic islands are few, and most of them serve the cell envelope. Of the 186 variable regions of eight or more genes, four recur at an equivalent position in nine genomes or more, and three of them encode lipopolysaccharide synthesis, namely the O-chain island (HVR2)^19^, a lipid A biosynthesis block, and a block encoding the KDO transferase and the lipid A exporter MsbA (Figure 3A).

Matching regions across genomes by the pair of core clusters flanking them, which makes the comparison independent of genome rearrangement (Figure S3A), showed that most variable positions are shared. Sixty-five positions are present in all fifteen genomes, and 187 in at least eight, and the latter hold 83% of all variable regions observed, whereas only 8.2% of regions lie at a position found in a single genome. The population therefore maintains a conserved set of chromosomal slots, each holding one or two genes and each occupied by a different sequence variant in different cells, alongside a small number of large strain-dependent islands.

At the 76 positions that exist in all fifteen genomes, that is, whose two flanking core clusters are adjacent in every one of them, and that are occupied in at least ten, the slot holds a gene of the same metaparalog family in 91.4% of cases, a different family in 6.5%, and is absent in 2.1%, with the two flanking core genes being directly contiguous. A third of the 174 apparently unrelated occupants still align with the majority family at ≥30% identity but over less than 70% of their length. Variations at these positions consist of one member of a metaparalog family replacing another.

Within these strain-dependent positions, occupancy is mosaic. Grouping at 90% amino acid similarity, the occupants of the 219 positions at which a metaparalog lies between the same two core clusters in at least ten genomes resolved a median of five variants per position. The pool of metaparalog variants does not saturate. A single genome carries 202 ± 10 of them, and the fifteen together 1,179, of which 804 (68%) occur in only one genome, and the fifteenth genome still adds 51 (Heaps γ = 0.65; Figure 2C). The fifteen strains therefore sample only a fraction of the variation present in the population.

**Figure 2.**
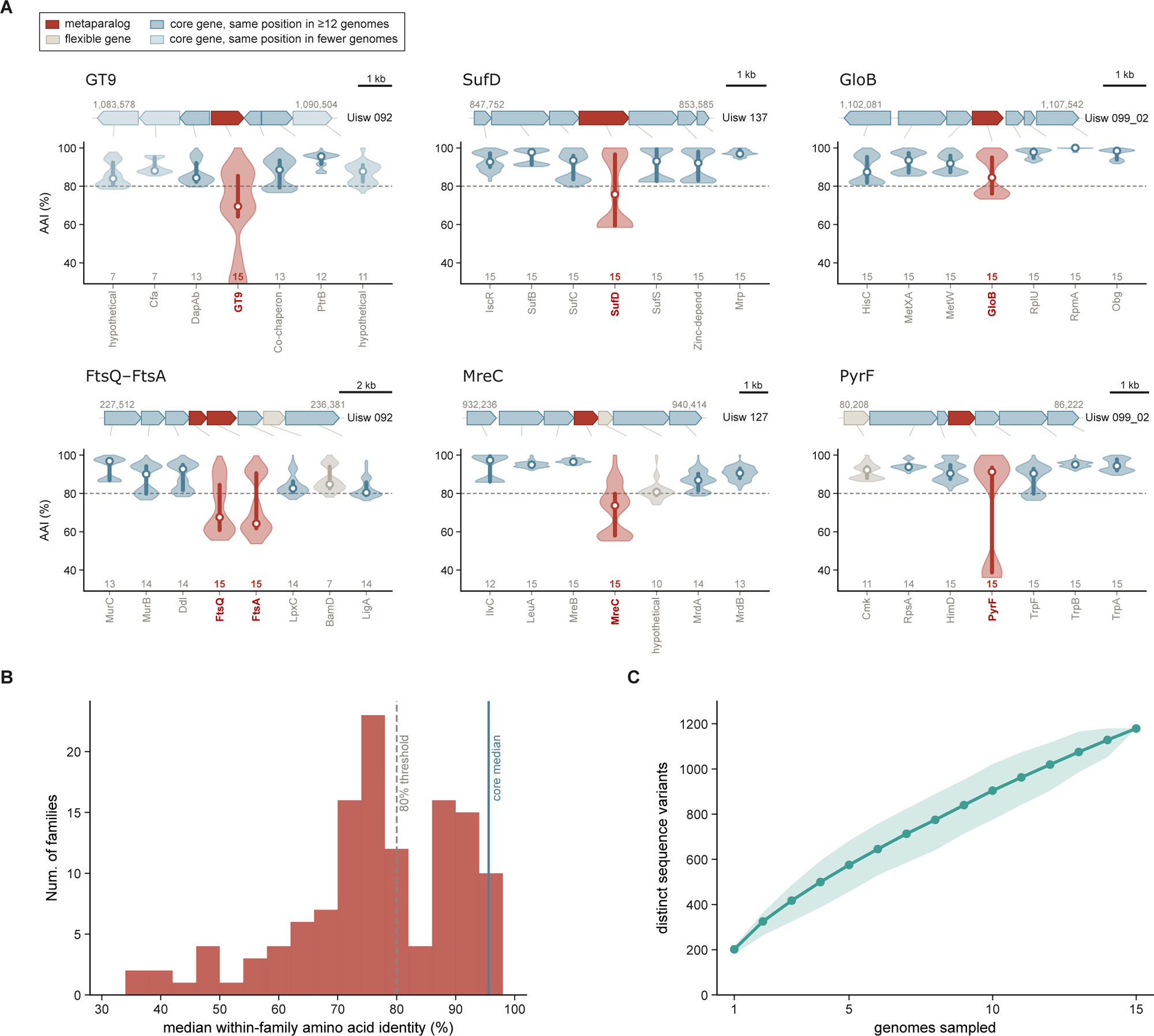
Sequence divergence of metaparalogs at conserved chromosomal positions. (**A)**, Six positions occupied by a single-copy metaparalog family in all fifteen genomes, each drawn to scale in a reference strain. Genes are coloured by pangenome status. Violins give the pairwise amino acid identity of each gene across the genomes carrying it (number of genomes below each violin); circle, median; bar, interquartile range; dashed line, 80% clustering threshold. (**B)**, Median within-family identity of the 127 single-copy metaparalog families; dashed line, 80% threshold. (**C)**, Accumulation of distinct variants (90% identity clusters) at the 219 positions occupied by a metaparalog in at least ten genomes, over 500 random orderings of the genomes (line, mean; shading, 5th–95th percentile).

A total of 127 metaparalog families, comprising 1,905 genes, are present as a single copy in all fifteen genomes and yet fall in the flexible fraction because their within-family identities are sharply split (Figure 2A,B). Fifty lie between 70 and 80%, 47 have a median above 80% and are flexible only because one or two strains carry a deeply divergent copy, and 30 fall below 70%, reaching 35%. For 43 of the 127 families, at least four neighbouring core genes occupy the same position in twelve or more genomes, providing a local reference. At these *loci*, the metaparalog was 78.5% identical between genomes at the median, compared with 93.3% for its flanking core genes, and it was the less conserved gene in 38 of the 43 *loci* (median difference 11.5 percentage points; two-sided Wilcoxon signed-rank test, *P* = 1.0 × 10⁻¹⁰). Among others, these *loci* include a GT9 heptosyltransferase of the lipopolysaccharide inner core, the Fe–S cluster scaffold SufD, the glyoxalase GloB, the divisome pair FtsQ–FtsA, and the elongasome component MreC (Figure 2A). A metaparalog and the core genes beside it sit in the same cells, and have therefore been inherited through the same lineages for the same length of time. The gap between 78.5% and 93.3% therefore reflects a property of the gene itself rather than of the lineage that carries it. The contrast is not universal and in five of the 43 loci the metaparalog is as conserved as its neighbours. PyrF, for instance, differs from its flanking core genes by only three percentage points and is flexible only because one variant falls below the clustering threshold.

Such divergence would be expected of mobile genetic elements, but as in gMED^9^, in the Arctic genomes, canonical mobility functions are almost absent. No transposase, insertion sequence, or conjugation system was detected in any of the fifteen, and the only site-specific recombinases present in more than one or two genomes are XerC and XerD, which resolve chromosome dimers; traces of past mobility are sporadic and confined to individual cells, phage-type integrases in Uisw_101 and Uisw_130, the latter beside a serine resolvase, an integrase fragment in Uisw_137, and a phage T5 Orf172 DNA-binding domain in Uisw_090 and Uisw_101. Unlike gMED, where eight islands were bounded by tRNA or tmRNA genes, the variable regions show no association with these integration hotspots. Of the 482 tRNA and tmRNA genes, 247 lie within 500 bp of a strain-variable region, close to the 255 ± 10 expected under random placement (*P* = 0.69).

### Strains carry different versions of the same envelope and transport functions

As in the Mediterranean gMED population,^9^ envelope biogenesis was the most enriched functional category among metaparalogs that is also represented in the core, accounting for 14.7% of them against 4.9% of core genes (odds ratio 3.39, q = 1 × 10⁻¹¹⁰; Figure S4A, Table S5c). The same bias emerged from signal peptide prediction, which relies on sequence alone and therefore also covers proteins without functional annotation. Signal peptides were predicted for 15.5% of metaparalog families and 5.2% of core clusters (odds ratio 2.18 after adjustment for protein length and annotation, P = 2.6 × 10⁻⁵), whereas proteins with transmembrane segments were not over-represented (odds ratio 0.95, P = 0.69; Table S5e). The bias therefore concerns proteins exported across the inner membrane. Because the genomes are complete, it could be traced to the machinery that builds the envelope. Twenty-two of the 127 families present as a single copy in all fifteen genomes carried a signal peptide, among them BamA, BamB and BamD of the β-barrel assembly machinery, the lipoprotein carrier LolA and the phospholipid transporter MlaA, and the three BAM components resolved into eleven, ten and eight distinct 80%-identity clusters across the fifteen genomes (Table S5b). Therefore, these results suggest that each cell assembles its envelope with its own version of a shared machinery.

COG composition varied only negligibly among genomes (χ^2^ = 369.0, d.f. = 252, P = 2 × 10^−6^; Cramér’s V = 0.062), so that strains differ in the sequences of their metaparalogs rather than in the functions these encode. The single appreciable departure was cell motility in Uisw_101 and Uisw_104 (Figure S4B, Table S5D). That departure reflects the one extracellular system that varies by presence and absence. Motility genes, absent from the core altogether, belong to a type IV pilus *locus* that occurred in nine genomes in two versions, one in five genomes and the other in four (Figure S5A). Pilin copy number varied more than ten-fold among carriers, from 2 in Uisw_116 and Uisw_121 to 25 in Uisw_101 and Uisw_104 (112 in total), whereas each assembly component was present once per genome, or twice for the PilT and PilU ATPases (Table S5F). The assembly components of the two versions form separate lineages, and each of the five components recovers the same split when analyzed separately (Figure S5B,F), indicating that the two systems have not exchanged parts.

Transporters, the other functional class that expands in the largest families (Figure 1E), show how this variation is organised along the chromosome. In the 22 transporter families occupying more than one syntenic position, members at the same position shared a median amino acid identity of 95.0% (n = 2,455 pairs) and members at different positions 32.5% (n = 2,808; Mann–Whitney P < 10^−300^ Table S5H). The difference was recovered in 20 of the 22 families (Wilcoxon signed-rank P = 1.6 × 10^−4^), and divergent members lay in unrelated genomic neighbourhoods (median Jaccard similarity of the six flanking genes, 0). The tricarboxylate transporters make this organisation explicit (Figure 3B,C). The 73 DctPQM operons of the population fell into 15 versions distributed over 13 chromosomal positions, and the 19 TctAB operons into five versions over eight positions. Operons of the same version were 97.2% (DctPQM) and 97.6% (TctAB) similar, compared with 32.3% and 40.0% for subunits at different positions, and at a given position the transporter subunits were at least as conserved as their flanking core genes (96.9% versus 94.2% for DctPQM, 98.2% versus 93.5% for TctAB; Table S5I-K). Divergence between versions is therefore found mostly between positions. A five-gene phosphate uptake block (genes for SphX, PstB, PhoU and two permeases) followed the same rule, occupying a single position in the five genomes that carried it, with 84–100% identity between copies (Table S5l).

**Figure 3.**
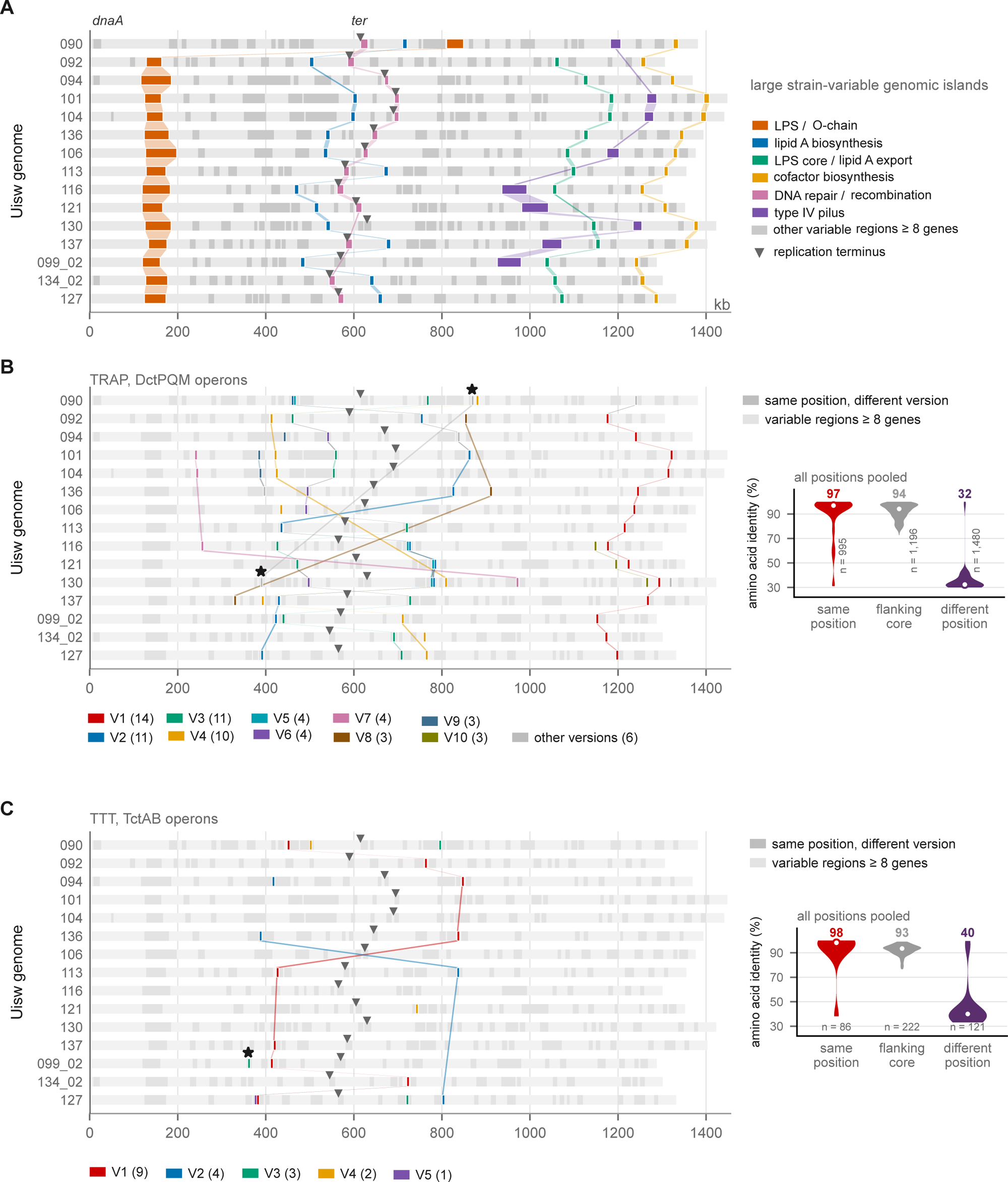
Chromosomal organisation of the variable genome and of its two tricarboxylate transport systems. **(A)**, Large variable regions in the fifteen Arctic genomes. Each chromosome is drawn as a horizontal bar beginning at *dnaA* and the triangle marks the replication terminus. Of the 186 variable regions of eight or more genes, those recurring at an equivalent syntenic position in nine genomes or more are coloured by function (four classes); the remainder are shown in grey. (**B)**, The DctPQM TRAP system. The 73 complete operons are mapped onto the same chromosomes and coloured by sequence version (V1–V15; a version being a set of operons whose three subunits are mutually at least 92% identical). The thirteen syntenic positions the system occupies are defined by the pair of core clusters flanking each operon and are therefore independent of coordinate. Shaded ribbons join operons of the same version in adjacent genomes. Stars mark position C00455|C00571, the only position carrying a TRAP operon in two genomes (Uisw_090, Uisw_130) and a TctAB operon in a third (Uisw_099_02). (**C)**, As (B) for the TctAB tricarboxylate system. 19 operons, five versions, eight positions. Right column, amino acid identity between operon subunits carried by different genomes (pairs at the same syntenic position, the six core genes flanking them, and pairs at different positions). The number of comparisons is given under each violin.

This diversification is confined to particular transporter classes and is older than the population itself. Tricarboxylate and phosphate transporters were absent from the core, and the three TRAP subunits were 6.7- to 8.4-fold enriched in the metaparalog pool, as was, to a lesser extent, the ZnuABC zinc uptake system. Potassium transporters and Na⁺/H⁺ antiporters, in contrast, were absent from the metaparalog pool, and magnesium transporters nearly so (2 metaparalogs against 14 core genes). ABC transporters, the most abundant family in both fractions, were distributed in proportion to the core (1.13-fold; Table S5G). The diversifying families were shared with the Mediterranean genomospecies. When clustered jointly with seven near-complete gMED genomes, 88–100% of their genes fell into families containing members from both oceans, yet identity between populations matched identity within each (DctP, 28.2% within gMED, within the Arctic and between them; TctA, 39.9%, 40.6% and 39.9%; Table S5M). The transporter variants of an Arctic cell are thus no more similar to one another than to those of a Mediterranean cell, indicating that each population samples a repertoire that diversified before the two genomospecies separated.

## DISCUSSION

### Metaparalog definition

Paralogous proteins are conventionally defined as arising by duplication, with subsequent divergence in the sequence space that leads to a shift in substrate specificity.^12,20^ In prokaryotes, however, new family members may more often arise by incomplete xenologous (sometimes called pseudoparalogs) replacement than by duplication.^11,21^ Acquisition has a major mechanistic advantage because two closely similar sequences coexisting in one genome are subject to gene conversion, which homogenizes them and erodes the divergence that functional differentiation requires. We have previously defined the concept of metaparalogs as homologous genes that carry out similar but diverging functions found within a local pangenome, that is, a population of cells belonging to the same species and retrieved from the same place and time (ideally from the same sample).

### Mechanistic aspects of metaparalog family generation

Following the framework established by Molina-Pardines et al.^9^ for Mediterranean gMED populations, we analyzed metaparalogs in the Arctic Ia.1 population to test whether this distributed paralog strategy occurs across different SAR11 species in contrasting environments. Having access to complete genomes has allowed us to delve into the mechanistic aspects that might be behind the generation of these vast pools of metaparalogs and, more generally, of the pangenome. Metaparalogs occur predominantly as single genes or small gene cassettes distributed throughout the chromosome. Describing them as “inserted” would, however, prejudge a mechanism that remains unresolved. When and how these variants became established is an elusive question. Their distribution provides little evidence that recent mobile-element-mediated transfer is the predominant mechanism, as only nine metaparalogs (0.1% of the total) occur within three genes of an annotated integrase, integrase fragment, or prophage-related element. Instead, their chromosomal positions are remarkably recurrent. A total of 187 variable *loci* shared by at least eight genomes account for 83% of the 3,055 variable regions identified, and these *loci* are defined by the same flanking core genes. This positional conservation is compatible with homologous replacement or recombination acting at pre-existing chromosomal sites. The finding that Arctic DctP variants are no more similar to each other than to Mediterranean variants (28.2% within each population, 28.2% between) needs to be emphasized. It means the metaparalog pool predates the speciation of Ia.1 and Ia.3/VII, which supports an ancient, conserved strategy rather than convergent evolution.

A related question is whether the high sequence diversity of these regions necessarily implies rapid turnover. Previous comparisons of O-chain biosynthetic gene clusters located in strain-variable HVR2^19^ indicated that their divergence can be comparable to that of the core genome itself, consistent with ancient rather than continuously renewed exchanges.^23^ The weak association of metaparalog sharing with ANI relationship is also indicative that their distribution predates the divergence of the strains. This issue is important because the frequently used term “hypervariable island” can be interpreted as implying unusually rapid evolutionary change, whereas a highly diverse *locus* may instead represent the long-term persistence of alternative alleles or gene cassettes.^24^ We therefore favour the more time-neutral term “strain-dependent island” for such regions.

### Evolutionary and ecological implications

We have already established that, as previously suggested^9,10^, markedly divergent metaparalogs (at least in the case of transporters) belong to divergent gene clusters containing different cognate metabolic pathways inferred to be involved in the metabolism of the transported molecule. The association of the elements of these gene clusters (transporters plus catabolism) might reflect the mobility of the whole cluster or the much slower process of gradual co-capture of two independently evolved components. Mechanistically, this could be due to MGE importing the whole assembly already pre-formed in another microbe or the slow evolution by the unlikely capture of the components. The analysis of the synteny carried out with our set of strains seems to indicate that each version of the gene cluster occupies a syntenically equivalent position in the genome so that the gene clusters are not recent acquisitions, they have evolved along with the genome remaining within the same domain or even region. Interestingly, nearly all the examples of location shift are in the form of a symmetric switch around the replication terminus (Figue 3B,C; for example, DctPQM V4 lies 11–19% of chromosome length from the terminus on either replichore) . This might be indicative of homologous recombination between the duplicating DNA strands as culprit of the rearrangement.

Regardless of how the diversity of metaparalogs is generated, their biological implications seem clear. They provide the population with an extensive range of metabolic and adaptive possibilities. From a kin selection perspective, the chances of ecological success are better, and the service to the community is much more complete (better ecosystem function). There is the question of the maintenance of such vastly diverse populations across time. That viruses fulfil an important role by preventing hoarding of the resources in a “kill-the-winner” dynamic is clearly illustrated and explained by the diversity of envelope components that provide each clone with identity markers for differential phage sensitivity.^25^ However, the fact that every cell has a different version of the outer-membrane protein assembly machinery (BamA/BamB/BamD), indicates a wholesale individualisation of the cell surface. That the population carries a large set of different envelope proteins might act as an additional benefit in itself beyond phage sensitivity. The outer membrane of a Gram-negative bacterium determines many properties of the cell as a whole, including its permeability to substrates or interaction with other cells (either directly or via cell vesicles). Finally, it is likely that even under quite restrictive conditions, clones could survive at semidormant state, avoiding extinction.

Our findings have broader implications for understanding bacterial adaptation in nutrient-limited environments. Paralog-mediated genomic flexibility may be a widespread strategy among streamlined marine bacteria, representing an alternative evolutionary solution to environmental complexity. Future research should investigate metaparalog patterns across additional marine bacterial groups and environments to test the generality of this adaptive mechanism.

### Limitations of the study

This study is based on a limited number of cultured isolates recovered from a single sample, and the relationships described here should be tested in larger collections of complete genomes spanning different locations, seasons and SAR11 lineages. Cultivation may not capture the full diversity of the natural population. Metaparalogs are defined by sequence-similarity thresholds, and their functions are inferred from sequence and genomic context rather than tested experimentally.

## Supporting information

Tabe S4

## RESOURCE AVAILABILITY

### Lead contact

Requests for further information and resources should be directed to Francisco Rodriguez-Valera

### Materials availability

#### Data and code availability

This study did not generate new unique reagents.

This paper analyzes existing, publicly available data. The 15 Arctic *Pelagibacter* genomes are available at NCBI under BioProject PRJNA1129510 (GenBank accessions CP159809.1–CP159824.1), and the gMED genomes as described in Molina-Pardines et al.^9^ The Prokka annotations of the fifteen genomes analyzed, the pangenome partition, the metaparalog families, all alignments and gene trees, and the source data for every figure have been deposited at Zenodo and are publicly available as of the date of publication at https://doi.org/10.5281/zenodo.23139235.

## ACKNOWLEDGMENTS

This work was supported by grant “SIMBAV” PID2024-162730NB-I00 from the Spanish Ministerio de Ciencia, Innovación y Universidades.

## AUTHOR CONTRIBUTIONS

Conceptualization, F.R.-V. and A.-B.M.-C.; methodology, A.-B.M.-C. and R.R.; writing F.R.-V. and A.-B.M.-C.; review & editing, A.-B.M.-C., F.R.-V. and R.R.

## DECLARATION OF GENERATIVE AI AND AI-ASSISTED TECHNOLOGIES

During the preparation of this work, the authors used Claude (Anthropic) and ChatGPT (OpenAI) to improve the language and readability of the manuscript, to check the consistency of the reported values, and to assist with writing and debugging analysis scripts.

## METHODS

### KEY RESOURCES TABLE

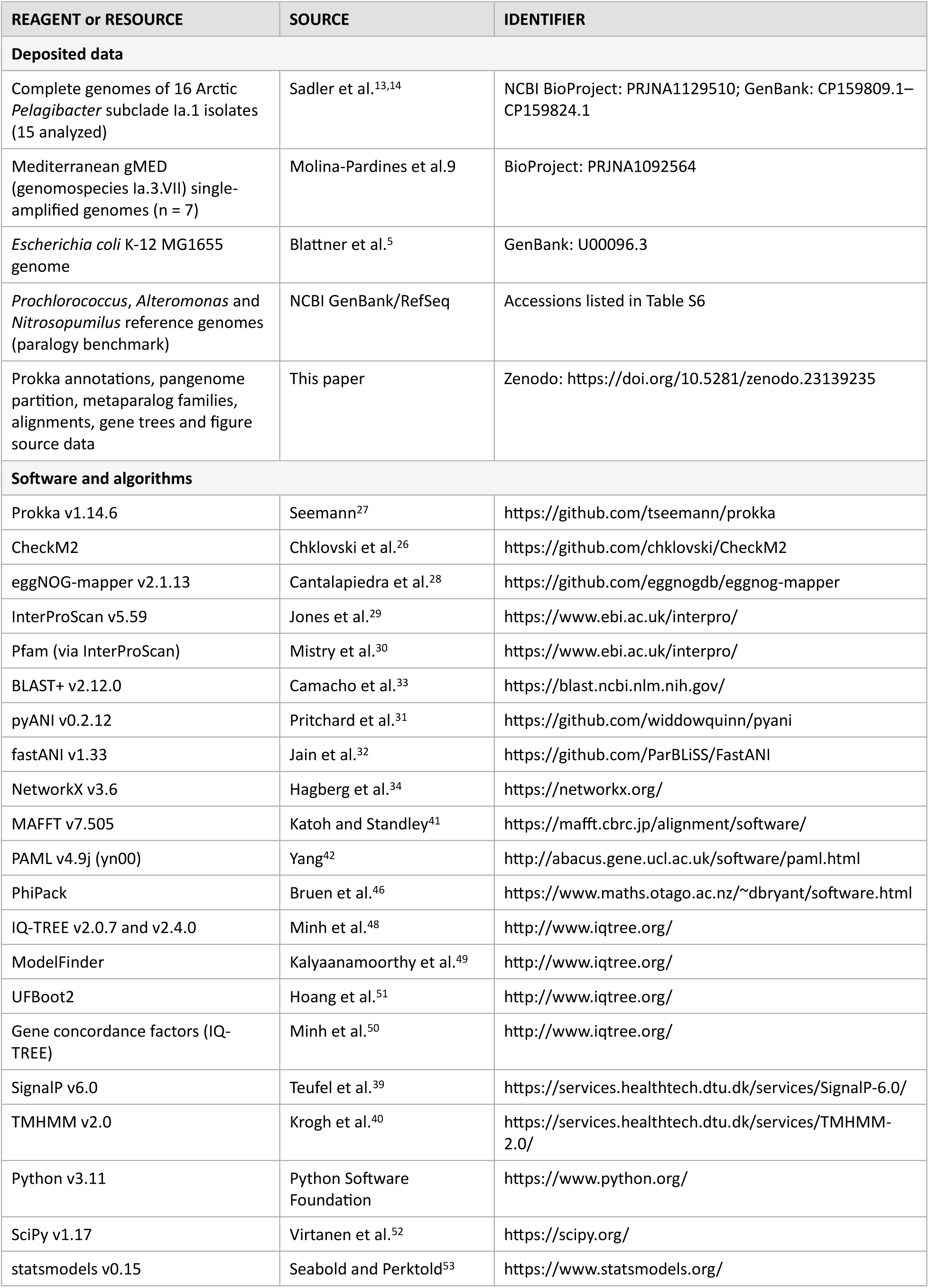

### Methods Details

#### Genome dataset, quality control and annotation

We initially analysed the 16 closed, circular *Pelagibacter* subclade Ia.1 genomes reported by Sadler et al.^13,14^ (Table S1). All strains were obtained by dilution-to-extinction cultivation from a single sub-ice seawater sample collected in Fram Strait (81.04° N, 10.62° E), 25 m below the ice–water interface, in May 2023. The original genomes were assembled from Oxford Nanopore reads with Flye v2.9.1 and polished with Medaka v1.7.2.^13,14^ Of the 16 genomes, Uisw_114 was excluded from subsequent analyses because it had the lowest sequencing coverage of the collection (11×) and, after uniform reannotation, was also an outlier in several independent quality metrics, with the lowest CheckM2^26^ completeness (92.8%), the shortest mean protein length (262 amino acids (aa) against 287-302 aa in the other genomes), and the largest number of predicted ORFs and proportion of hypothetical proteins (Tables S1 and S2). These features indicate residual gene fragmentation relative to the rest of the collection. The 15 genomes retained had CheckM2 completeness of 96.25–99.95% (mean 99.0%) and contamination ≤1.47% (Table S2). The seven most complete gMED genomes (89.5-99.1% estimated completeness) were obtained from Molina-Pardines et al.^9^

All chromosomes were oriented to begin at *dnaA* and reannotated with Prokka v1.14.6 under identical parameters.^27^ Functional annotation used eggNOG-mapper v2.1.13,^28^ from which COG categories were taken, and InterProScan v5.59,^29^ from which Pfam domains^30^ were taken. Mobile genetic elements were searched by annotations (transposases, insertion sequences, integrases, resolvases, conjugation systems, and prophage genes), and candidates were inspected individually. The XerC and XerD recombinases, which resolve chromosome dimers, were not counted.

#### Genome relatedness

Pairwise average nucleotide identity (ANI) values between the genomes were taken from Sadler et al.,^14^ who computed them with pyANI v0.2.12,^31^ and were reproduced with fastANI v1.33^32^ using default parameters, averaging the two reciprocal estimates of each pair. The relationship between ANI and the number of metaparalog families shared by each of the 105 genome pairs was assessed by Pearson correlation and by partial correlation controlling for the summed flexome size of the two genomes.

For each pair of genomes, conserved metaparalogs (mp) were the flexible genes of one genome with a homolog in the other under the metaparalog criterion, averaged over the two directions.

#### Pangenome partition

All proteins were compared with each other by BLASTP (BLAST+ v2.12.0; E ≤ 10⁻⁵)^33^. Two proteins were linked when they shared ≥80% amino acid identity over ≥70% of the length of both sequences, and clusters were defined as the connected components of the resulting graph (single linkage; NetworkX v3.6)^34^. A cluster was classified as core when present in at least 14 of the 15 genomes and as flexible otherwise. The 80% threshold matches the identity threshold used to cluster the Mediterranean gMED flexible genome,^9^ and the requirement for coverage of both sequences prevents linkage through isolated shared domains. The robustness of the partition was assessed for clustering thresholds of 70, 75, 80, 85, 90, and 95% identity and core-prevalence thresholds from 6 to 15 genomes (48 combinations; Table S3).

#### Metaparalogs and intragenomic paralogs

Homology was defined on the same BLASTP search as ≥30% amino acid identity over ≥70% of both sequences, with E ≤ 10⁻⁵, a threshold near the limit of reliable homology detection.^35^ A flexible gene was classified as a metaparalog when it had at least one such homolog encoded in a different genome of the population, and as an intragenomic paralog when the homolog was encoded in the same genome. Metaparalog families were defined as the connected components of the flexible proteins under the same criterion, and families of at least two proteins were analysed. Sensitivity to the identity threshold was evaluated at ≥40% (Supplementary Information).

To place intragenomic paralogy in context, the same procedure was applied to closed genomes of streamlined and non-streamlined bacterial and archaeal lineages, reannotated with Prokka under the same parameters, with *Escherichia coli* K-12 MG1655 as reference.^5^

#### Chromosomal position of metaparalogs and syntenic context

A variable region was defined as a maximal run of consecutive flexible genes bounded by core genes, taking chromosome circularity into account. The syntenic position of a variable region, or of any gene within it, was defined by the pair of core clusters flanking it, so that regions in different genomes bounded by the same pair were assigned to the same position irrespective of their content, orientation, or chromosomal coordinate.

The distribution of metaparalogs along the chromosome was expressed in windows of 2% of chromosome length and tested against uniformity in each genome with a Kolmogorov–Smirnov test, with Benjamini–Hochberg correction across genomes.^36^ Replication termini were located at the maximum of the cumulative GC skew computed in 5-kb windows.^37^ Consecutive genomes were aligned with BLASTN (BLAST+ v2.12.0; E ≤ 10⁻⁵), retaining non-overlapping blocks ≥500 bp at ≥70% identity, and a block was scored as inverted when its orientation differed between the two genomes. The proximity of variable regions to tRNA and tmRNA genes (within 500 bp) was tested against 1,000 random placements.

The sequence variants occupying each syntenic position with a metaparalog in at least ten genomes were clustered at 90% amino acid identity by single linkage on the same BLASTP search, and their accumulation was computed over 500 random orders of the genomes and fitted to Heaps’ law by least squares on log-transformed values.^38^ For metaparalog families present as a single copy in all 15 genomes, positions where at least four neighbouring core genes occupied the same position in at least 12 genomes were used to compare the between-genome identity of the metaparalog with that of its flanking core genes (two-sided Wilcoxon signed-rank test across *loci*).

#### Functional classification

COG categories of metaparalogs and core genes were compared with two-sided Fisher’s exact tests with Benjamini–Hochberg correction, counting genes with at least one COG assignment and counting a gene in each of its categories. Heterogeneity among genomes was tested with a χ^2^ test of independence on the genome-by-category table of the categories assigned to at least 15 metaparalogs, and departures were expressed as standardised residuals and summarised by Cramér’s V. For Figure 1, each protein was assigned hierarchically to transport (transporter Pfam domain or annotation), cell envelope (COG M), metabolism (COG C, E, F, G, H, I, P or Q), information and regulation (COG J, A, K, L, B, D, Y, V, T, U or O), hypothetical or unknown (no annotation, or COG S or R), or other.

Transporter classes were defined by Pfam domains, and the enrichment of each class was expressed as the ratio of its frequency among metaparalogs to its frequency in the core. Transporter families were defined as metaparalog families of at least six proteins in which at least 30% of the members carry a transporter Pfam domain. For each family, the identities between members encoded in different genomes, taken from the BLASTP search, were compared between members at the same and at different syntenic positions. The genomic neighbourhoods of the positions occupied by the same family were compared by the Jaccard index of the eggNOG orthologous groups of the three genes on either side.

An operon was scored as complete when all of its subunits were encoded within six consecutive genes, DctP (PF03480), DctQ (PF04290) and DctM (PF06808) for the TRAP system, and TctA (PF01970) and TctB (PF07331) for the tripartite tricarboxylate system. Overlapping calls in the same genome were merged, keeping the call with the longest DctP or TctA. The identity of two operons was the mean of the identities of their corresponding subunits, and operons were grouped into versions by single linkage at a mean identity of 80%. The syntenic position of each operon was defined by its flanking core clusters.

The seven gMED genomes were reannotated with Prokka v1.14.6 under the same parameters, and transporter subunits were identified by the same Pfam criteria. Proteins of both populations (Arctic and Mediterranean) were pooled and clustered with the metaparalog criterion (≥30% identity over ≥70% of both sequences, E ≤ 10⁻⁵, single linkage), and pairwise identities within each population were compared with those between populations.

Signal peptides were predicted with SignalP v6.0^39^ and transmembrane helices with TMHMM v2.0.^40^ Because a signal peptide is frequently reported by TMHMM as an N-terminal helix, only helices beginning after residue 60 were counted. To avoid pseudoreplication, the unit of analysis was the metaparalog family, the core cluster or the remaining flexible 80% cluster, scored positive when more than half of its members were. Enrichment relative to the core was tested by logistic regression with the logarithm of the median protein length and hypothetical annotation as covariates.

Type IV pilus components were identified by their InterPro entries. In Uisw_116, PilC and the PilM–PilN gene are each split into two consecutive open reading frames, which were joined. Each assembly component (PilQ, PilC, PilT, PilD and PilM–PilN) was aligned with MAFFT v7.505 (L-INS-i), and maximum-likelihood trees were inferred with IQ-TREE v2.4.0, for each component separately and for their concatenation, with the model selected by ModelFinder and 1,000 ultrafast bootstrap replicates.

#### Evolutionary-rate and selection analyses

Coding sequences were extracted from the GenBank files and checked against their Prokka translations. Proteins were aligned with MAFFT v7.505 (L-INS-i)^41^ and back-translated to codon alignments. Pairwise non-synonymous (dN) and synonymous (dS) distances were estimated with yn00 (PAML v4.9j)^42^ by the methods of Nei and Gojobori^43^ and Yang and Nielsen,^44^ after removing codons with a gap in either sequence of the pair.

All pairs of orthologues from the 690 core clusters present as a single copy in all 15 genomes were compared with all the pairs of metaparalogs of the same family occupying the same syntenic position in different genomes (959 position–family groups), following the comparison of core and non-core genes within a single species by Bohlin et al.^45^ Pairs with dS = 0 (where dN/dS is undefined) and pairs with dS ≥ 1 (where synonymous sites approach saturation) were excluded. The two sets were compared with two-sided Mann–Whitney tests on pairs and on per-*locus* medians. As an independent check, reciprocal best hits between the metaparalogs of every pair of genomes were taken from the BLASTP search, aligned individually (global alignment, BLOSUM62), back-translated, and retained when they shared ≥100 aligned codons and 0.01 ≤ dS ≤ 2. Yang–Nielsen estimates were considered stable when 0.01 ≤ dS ≤ 2 and dN/dS was finite.

#### Recombination and gene-tree concordance

Recombination was tested with PhiPack, using the pairwise homoplasy index (PHI)^46^ and the neighbour similarity score (NSS)^47^, each against 1,000 permutations of site order with the default window. Tests were applied to the codon alignments of the 690 single-copy core clusters and of 508 metaparalog *loci*, defined as position–family groups present as a single copy in at least four genomes (genomes with more than one copy at the position were omitted). An alignment was scored as recombinant when PHI and NSS were both significant at P ≤ 0.05. Proportions were compared with Fisher’s exact test and, to account for differences in statistical power, with a logistic regression on the number of sequences and the logarithm of the number of informative sites.

A maximum-likelihood tree was inferred for each locus with IQ-TREE v2.0.7,^48^ the substitution model being selected by ModelFinder.^49^ Gene concordance factors^50^ were calculated with IQ-TREE on the 12 internal branches of a reference phylogeny inferred with IQ-TREE v2.4.0 from a 624,847-nt supermatrix of 690 core genes (GTR+F+G4, 1,000 ultrafast bootstrap replicates)^51^. Core and metaparalog concordance factors were compared across branches with a two-sided Wilcoxon signed-rank test.

## QUANTIFICATION AND STATISTICAL ANALYSES

Analyses were performed in Python v3.11 with SciPy v1.17,^52^ statsmodels v0.15^53^ and NetworkX v3.6.^34^ All tests were two-sided. Independent distributions were compared with Mann–Whitney U tests, paired values with Wilcoxon signed-rank tests, and proportions with Fisher’s exact tests. Multiple tests within a family were corrected by the Benjamini–Hochberg procedure. Correlations were Pearson or Spearman as stated. The statistical test, the value and meaning of n, and the dispersion measures are given in the Results and figure legends. Permutation tests used fixed random seeds, so that every value can be reproduced from the deposited code.

## SUPPLEMENTAL FIGURES

**Figure S1. Organization of the O-chain biosynthesis island in the fifteen Arctic Pelagibacter genomes, related to Figure 1.** Each island is drawn to scale between its flanking core genes. Genes are colored by predicted role (glycosyltransferases; sugar synthesis and modification; export and polymerization; other annotated functions; hypothetical proteins), and the rRNA operon (16S–tRNA-Ile–tRNA-Ala–23S) that bounds the island in fourteen genomes is shown in purple. Chromosomal coordinates of each island, measured from the start of *dnaA*, are given below the genome name, and the number of genes at right. Gray ribbons join reciprocal best BLASTP hits between proteins of adjacent genomes (≥30% amino acid identity over ≥70% of both sequences), shaded by amino acid identity; ribbons between rRNA genes are shaded by nucleotide identity. In Uisw_090 the island lies at a different chromosomal position, between *garD* (coding for a (2R)-sulfolactate sulfo-lyase subunit beta) and a gene for a TauE-like transmembrane protein.

**Figure S2. Selection and recombination in core genes and metaparalogs. (A),** Pairwise dN/dS (Nei–Gojobori) for every pair of single-copy core orthologues (690 clusters) and for every pair of metaparalogs of the same family occupying the same syntenic position in different genomes (959 position–family groups), of which 840 contributed pairs with 0 < dS < 1. Violins show the distribution on a logarithmic scale; the point marks the median (given above each violin) and the bar the interquartile range; the dashed line marks dN/dS = 1 (Mann–Whitney P < 10⁻³⁰⁰). **(B),** Non-synonymous against synonymous distance for reciprocal best hits between the metaparalogs of every pair of genomes, with at least 100 aligned codons and 0.01 ≤ dS ≤ 2 (pairs with dN = 0 are not drawn). The dashed line marks dN = dS. **(C),** Yang–Nielsen against Nei–Gojobori dN/dS for the pairs in (B) with stable Yang–Nielsen estimates (0.01 ≤ dS ≤ 2 and finite dN/dS). **(D),** Percentage of alignments with significant homoplasy (P ≤ 0.05; 1,000 permutations of site order, PhiPack) by the pairwise homoplasy index (PHI), the neighbour similarity score (NSS) and both, for the 690 single-copy core clusters and 501 metaparalog *loci* (position–family groups present as a single copy in at least four genomes). Bars, Wilson 95% confidence intervals. The adjusted odds ratio is from a logistic regression on the number of genomes and the logarithm of the number of informative sites. **(E),** Gene concordance factor of each internal branch of the maximum-likelihood reference phylogeny, for core and metaparalog gene trees (IQ-TREE), joined by a line. Branches are named by the genomes on the smaller side of the bipartition and ordered by their core concordance factor; the ultrafast bootstrap support of each branch in the reference tree is given at right, in bold when below 95.

**Figure S3. Metaparalogs distribution across the chromosomes in all fifteen Arctic *Pelagibacter* genomes. (A),** Stacked whole-genome alignment. Chromosomes are drawn to scale and oriented at *dnaA*. Genomes are ordered to maximise synteny between neighbouring pairs. Ribbons join regions aligned by BLASTN between neighbouring genomes and are shaded by nucleotide identity. Values between genomes give the mean identity of the aligned blocks and the number of inverted blocks out of the total. Vertical lines within each chromosome mark the position of every metaparalog, coloured by the functional class assigned from its InterPro and Pfam domains (ABC transporters (PF00005, PF00664, PF04069), TRAP transporters (PF03480, PF04290, PF06808), TTT transporters (PF01970, PF03401, PF07331), type IV pilus components (PF07963, PF05157, PF00482, PF11104, PF05137) and ammonium transporters (PF00909); all remaining metaparalogs are shown in grey). Black triangles indicate the replication terminus. **(B),** Density of metaparalogs along the chromosome, expressed as the percentage of each genome’s metaparalogs falling in successive windows of 2% of chromosome length. Thin grey lines, individual genomes (437–595 metaparalogs each), each normalised to its own total; black line, mean of the fifteen genomes; dashed line, the 2% expected under a uniform distribution; shaded band, range of terminus positions. Departures from uniformity were not significant in any of the fifteen genomes after correction for multiple testing (Kolmogorov–Smirnov D = 0.042–0.077, Benjamini–Hochberg q > 0.05), and the mean profile across the population is indistinguishable from uniform (χ^2^ = 8.13, d.f. = 49, P > 0.999).

**Figure S4. Functional composition of the metaparalog pool and its distribution across genomes. (A),** Enrichment or depletion of each category among metaparalogs relative to the core, as the log₂ ratio of their relative frequencies. Significance is from two-sided Fisher exact tests corrected across categories by the Benjamini–Hochberg procedure (*** q < 10⁻¹⁰, ** q < 10⁻⁵, * q < 0.05). **(B),** Bipartite network linking the fifteen genomes (outer circle, blue) to COG categories (inner circle, orange squares, sized by the total number of metaparalogs assigned to that category. Genomes are arranged by ANI (italic values between adjacent genomes). Edge width is proportional to the percentage of that genome’s metaparalogs assigned to the category (below 1% are omitted). Edges are coloured by the standardised residual of a χ^2^ test of independence across all 285 genome-category combinations (red, over-represented (z > 2); blue, under-represented (z < −2); grey, as expected). Categories assigned to fewer than 15 metaparalogs in total were excluded from the test. *COG categories: C, energy production and conversion; D, cell cycle control and division; E, amino acid transport and metabolism; F, nucleotide transport and metabolism; G, carbohydrate transport and metabolism; H, coenzyme transport and metabolism; I, lipid transport and metabolism; J, translation and ribosome biogenesis; K, transcription; L, replication, recombination and repair; M, cell wall, membrane and envelope biogenesis; N, cell motility; O, post-translational modification, protein turnover and chaperones; P, inorganic ion transport and metabolism; Q, secondary metabolite biosynthesis, transport and catabolism; S, function unknown; T, signal transduction; U, intracellular trafficking and secretion; V, defence mechanisms*.

**Figure S5. The nine pilus-carrying genomes encode two independently acquired type IV pilus systems that differ in architecture and genomic context. (A)**, Organisation of the pilus *locus* in the nine carrier genomes. Genes are drawn to scale in chromosomal coordinates and aligned on the start of *PilC*; arrowheads indicate the direction of transcription. Roles were assigned from InterPro entries; pilin (PF07963/IPR012902), PilC/GspF platform, PilQ secretin, PilT ATPase, PilD prepilin peptidase, the PilM–PilN fusion protein, the PilY1 tip adhesin, type Va autotransporters (PF03797) and the anti-σ factor antagonist with its cognate RsbW kinase; flanking genes are coloured by pangenome partition. First and last chromosomal coordinates of each interval are given below it, measured from the start of *dnaA*, at which all genomes are oriented. The maximum-likelihood tree on the left is inferred from the concatenated assembly machinery (PilQ, PilC, PilT, PilD and PilM–PilN; 1,550 aligned positions) and orders the rows; branches are coloured by system and node labels are ultrafast bootstrap support from 1,000 replicates, with values below 70 not shown. The total number of pilin genes encoded by each genome, including copies outside this locus, is given at right. No component of either system belongs to the core genome, and the six remaining genomes of the population, listed below, encode no pilin, no assembly component, and no pilus locus at all. (B-F), Maximum-likelihood phylogenies of each assembly component analysed separately, with its own scale bar in substitutions per site. All five recover the same bipartition with maximal support, indicating that the two systems have not exchanged components. Trees are unrooted and drawn from Uisw_090.

In Uisw_116, *PilC* and the *PilM–PilN* gene are each annotated as two consecutive open reading frames and were joined for the phylogenetic analyses (see Methods).

## SUPPLEMENTAL TABLES

**Table S1. Genomic features of the 16 Arctic *Pelagibacter* subclade Ia.1 genomes, related to Figure 1.** Accession, genome size, GC content, numbers of CDS and RNA genes, coding density, protein length, and fraction of hypothetical and COG-annotated proteins. Uisw_114, excluded from the analyses, is shown for comparison.

**Table S2. CheckM2 completeness and contamination of the Arctic Pelagibacter genomes, related to Figure 1.**

**Table S3. Sensitivity of the pangenome partition and of the metaparalog fraction to clustering identity and core-prevalence thresholds, related to Figure 1.** Forty-eight combinations of clustering identity (70–95%) and core prevalence (6–15 of 15 genomes). The partition used throughout (80% identity, ≥14 of 15 genomes) is highlighted.

**Table S4. Selection and recombination in core genes and metaparalogs, related to Figure 2.** (A) Summary statistics. (B) Pairwise dN and dS for single-copy core orthologs and for metaparalogs of the same family at the same syntenic position. (C) PHI, NSS and MaxChi tests per *locus*. (D) Gene concordance factors for each internal branch of the core-genome phylogeny. (E) Composition of each *locus*.

**Table S5. Functional classification of the metaparalog pool, related to Figures 1 and 3.** (A) Annotation of the 7,424 metaparalogs. (B) The 897 metaparalog families. (C) COG enrichment relative to the core. (D) COG composition per genome. (E) Signal peptides and transmembrane helices. (F) Type IV pilus components. (G) Transporter classes. (H) Transporter families and identity within and between syntenic positions. (I–K) DctPQM and TctAB operons, versions and within-position identity. (L) The phosphate uptake block. (M) Comparison with the Mediterranean gMED genomes.

