## Supplementary figures and images for "Redundancy relocated: the distributed metaparalog pool of an Arctic *Pelagibacter*ales population"

### Figure S1

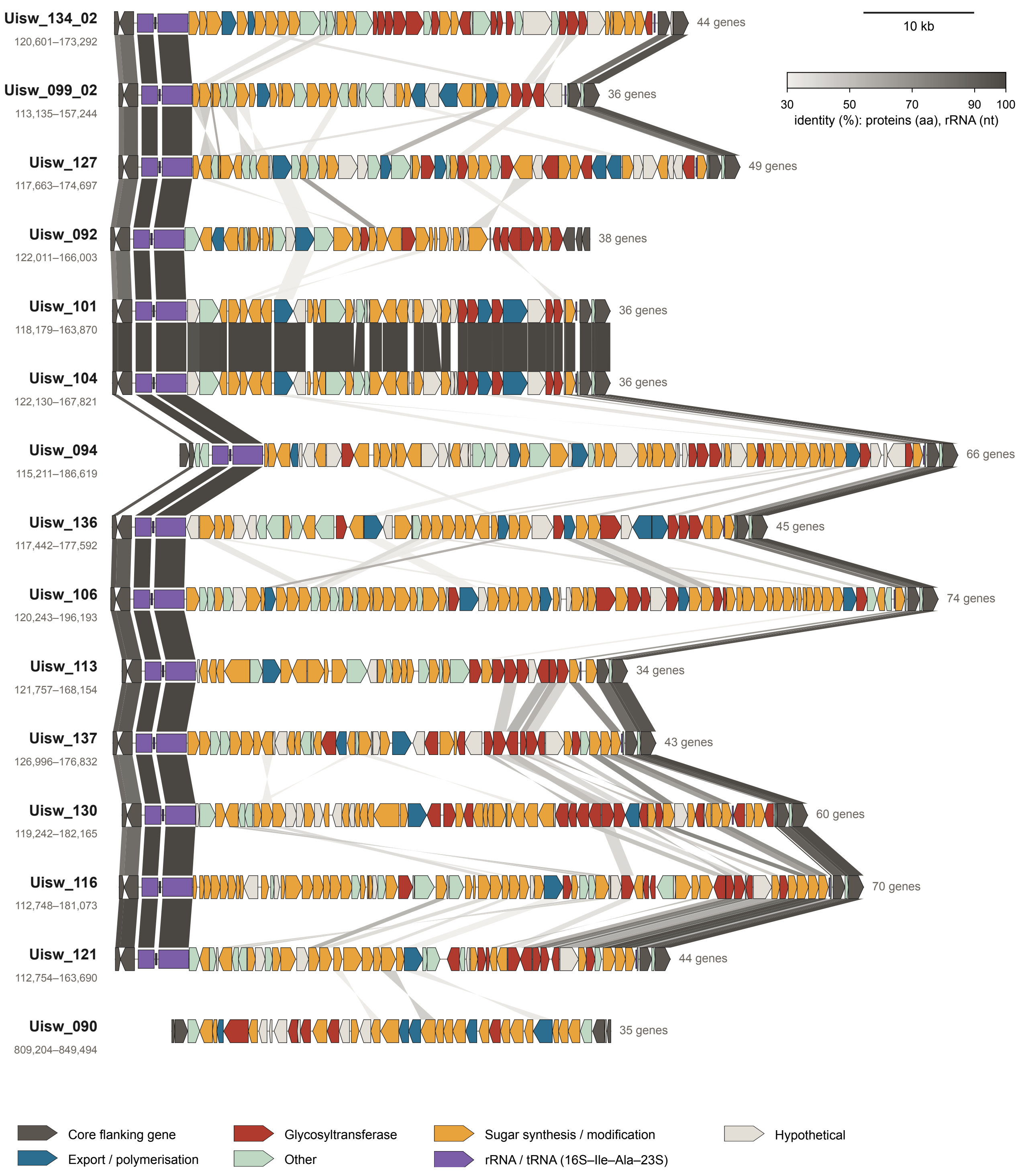

### Figure S2

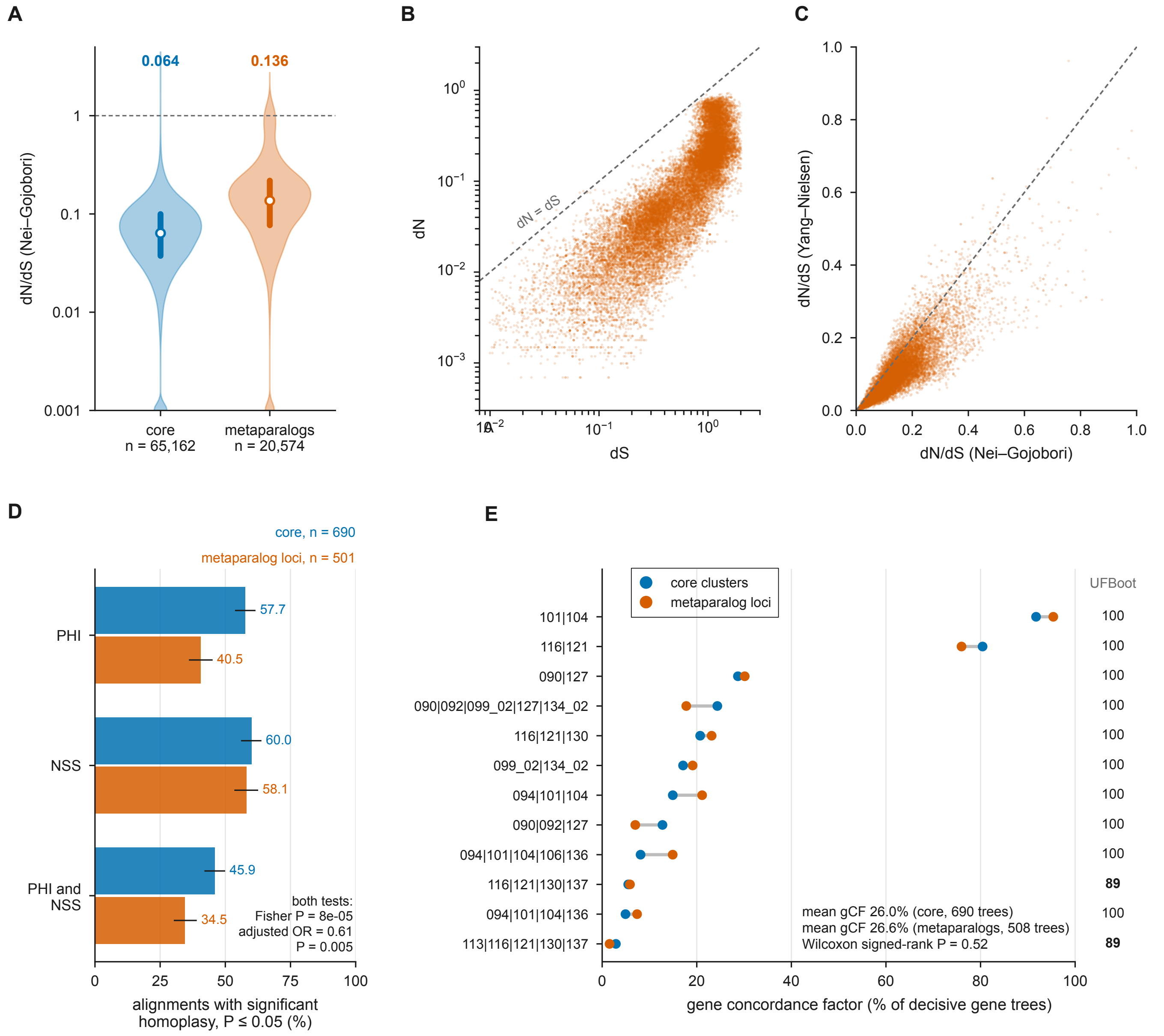

### Figure S3

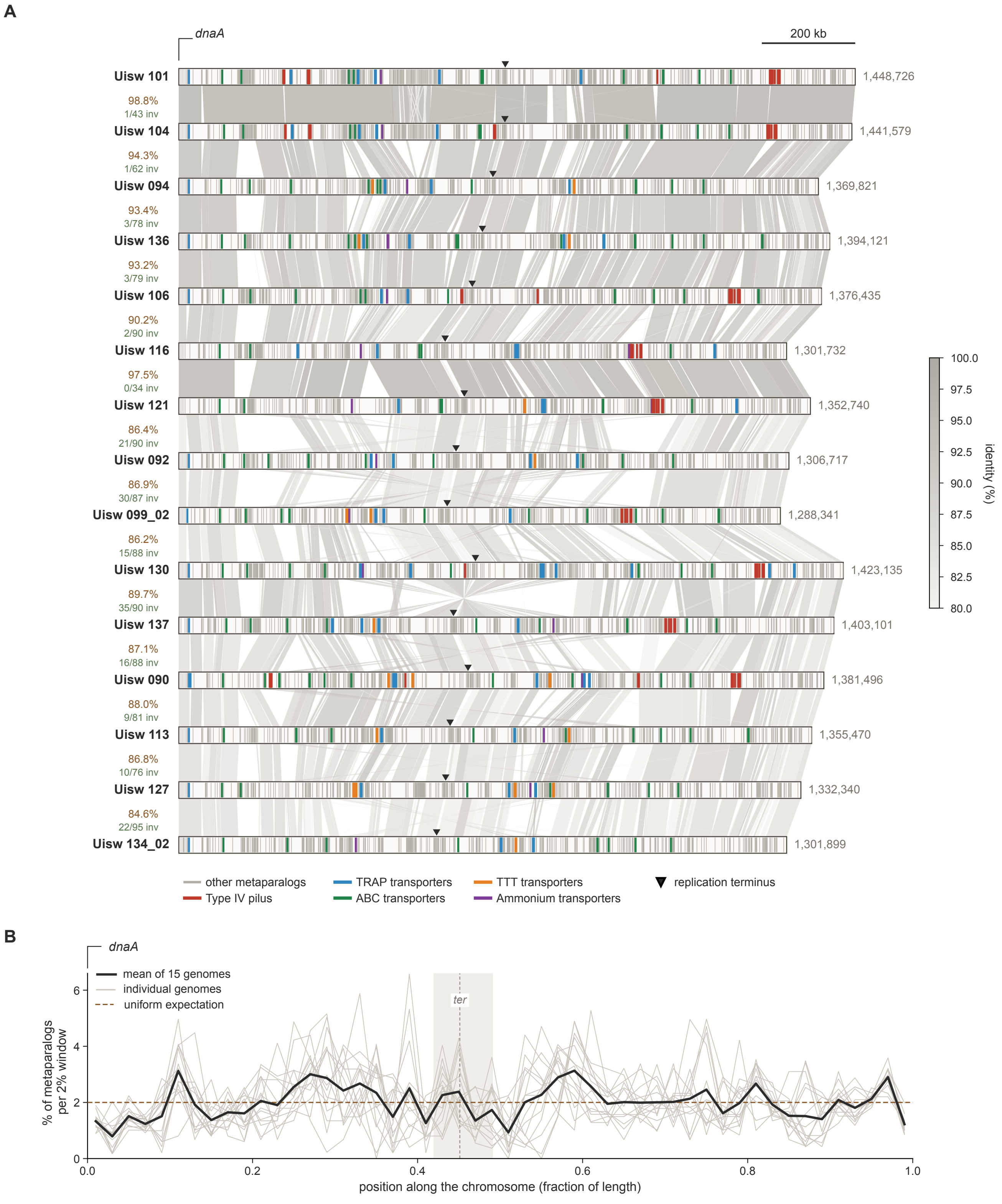

### Figure S4

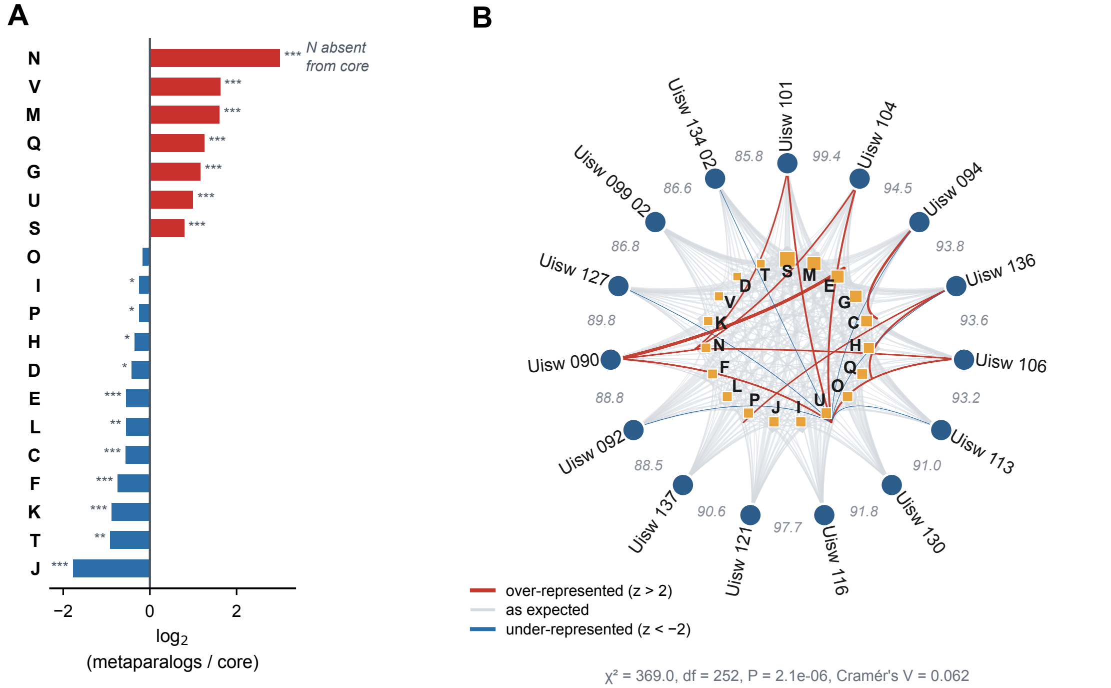

### Figure S5

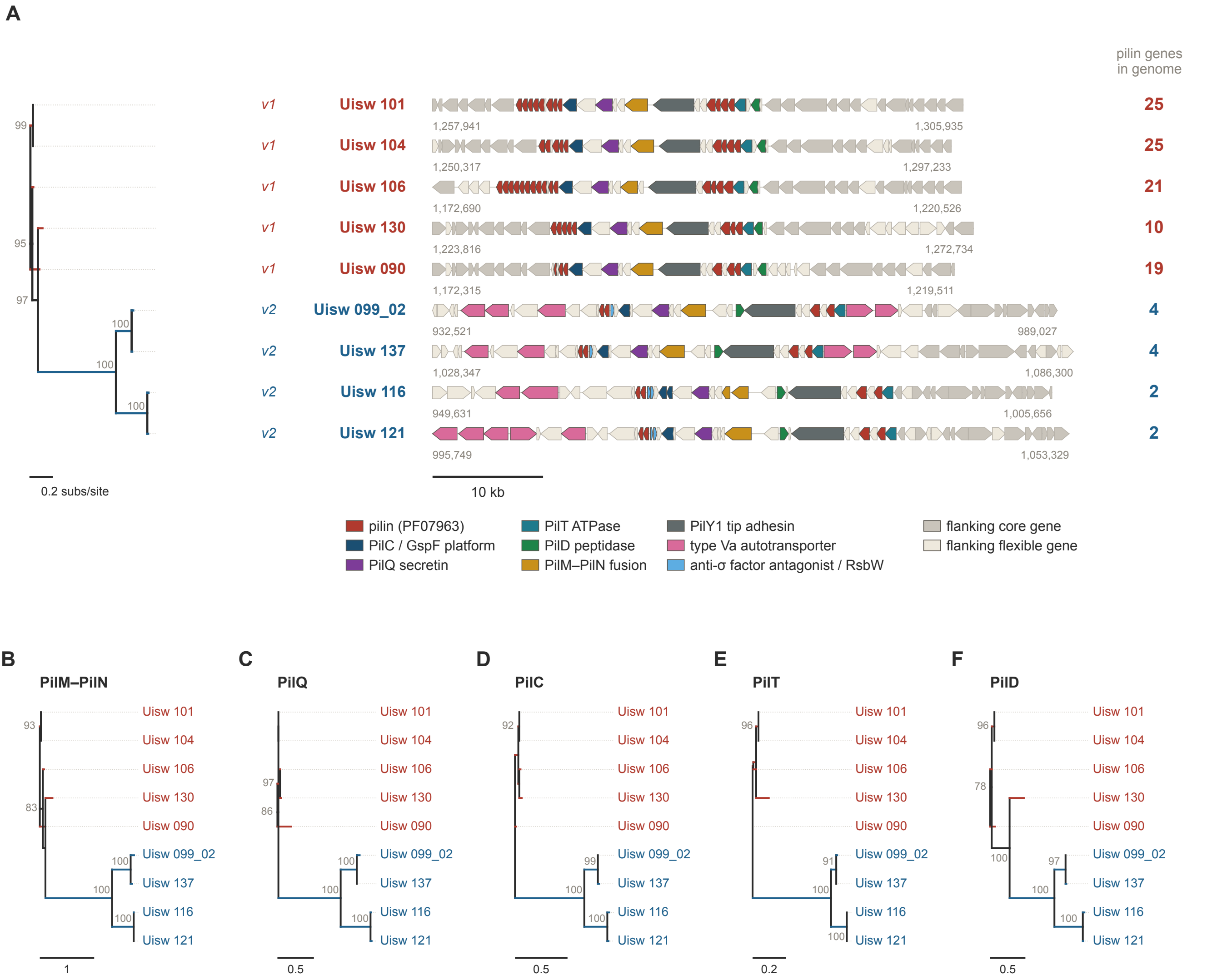
