## Supplementary Information for "Redundancy relocated: the distributed metaparalog pool of an Arctic *Pelagibacter*ales population"

**Paralogy scales with proteome size across bacterial lineages**

In *Pelagibacter* and other streamlined aquatic heterotrophs, intragenomic paralogs are far rarer than in *E. coli*. This has been read as straightforward evidence that streamlined lineages have traded innovative capacity for efficiency. Across the 15 closed Arctic genomes, 11.0 ± 1.2% of genes had at least one paralog elsewhere in the same genome (2,420 of 21,964; range 8.5–12.6%). Mobile elements were essentially absent (0.4 per genome; six proteins in total) and excluding them changed the estimate by less than 0.02 percentage points. Hence, the low paralog content is not an artefact of mobile element annotation. This is the lowest value of any lineage examined: under identical criteria, *E. coli* K-12 reached 27.8%, a 2.5-fold difference.

Paralog content increased monotonically with proteome size across the bacterial lineages: 11% in *Pelagibacter* (1,464 proteins per genome), 13.1% in *Prochlorococcus* (2,072), 16.4% in *Alteromonas mediterranea* (3,815), 17.4% in *Alteromonas macleodii* (3,946) and 27.8% in *E. coli* (4,305). Across the 58 individual bacterial genomes, the correlation was strong (r = 0.809, r² = 0.655, p = 1.5 × 10⁻¹⁴; 2.75 percentage points per 1,000 proteins) and held within genera (*Prochlorococcus* alone, r = 0.718). Raising the identity threshold to 40% reduced all estimates but preserved the ordering (Supplementary Information Figure 1, Supplementary Information Table 1).

The excess of *E. coli* over *Alteromonas*, a lineage of essentially identical genome size, is not explained by mobile elements: *Alteromonas* genomes carry 34–36 transposase-like proteins on average and their removal lowers the estimate by only 0.2–0.4 points. The largest *E. coli* paralog family (51 members) comprises ATP-binding components of ABC transporters, and only one of the eight largest families consisted of mobile elements, consistent with earlier observations that large gene families in bacteria are enriched in transporters (Pushker et al. 2004).

Within *Prochlorococcus*, the 26 genomes separated into two groups that track ecotype. High-light-adapted strains and the equally reduced low-light LLII strain SS120 clustered at 1,833–1,982 proteins and 10.1–12.9% paralogy (n = 14), whereas low-light LLI strains such as NATL1A, NATL2A and MIT0917 clustered at 2,108–2,365 proteins and 13.9–19.1% (n = 10). The two LLIV strains departed from this pattern: MIT9313 and MIT9303 carry the largest proteomes of the genus (2,552 and 2,728 proteins) yet reached only 13.8% and 14.6%, below the LLI strains, which carry 190–620 fewer proteins.

*Nitrosopumilus* isolates departed more sharply, reaching 18.4 ± 1.9% with a proteome size almost identical to that of *Prochlorococcus* (2,062 against 2,072 proteins), and were excluded from the regression on that basis. Whether this reflects a distinct history of gene duplication in Thaumarchaeota or an annotation artefact remains open: Prokka left 61–76% of *Nitrosopumilus* proteins as hypothetical, against 40–47% in the bacterial genomes.

Genome size therefore does not account for paralog content within the genus, and the expansion seen in LLI appears to be a lineage-specific feature rather than a simple consequence of a larger genome. Across lineages, however, the trend holds, and the low paralog content of *Pelagibacter* is close to what its proteome size predicts rather than a distinctive loss: streamlined genomes carry few paralogs largely because they carry few genes. What distinguishes this population is not the absence of redundancy but its location, outside the individual genome.

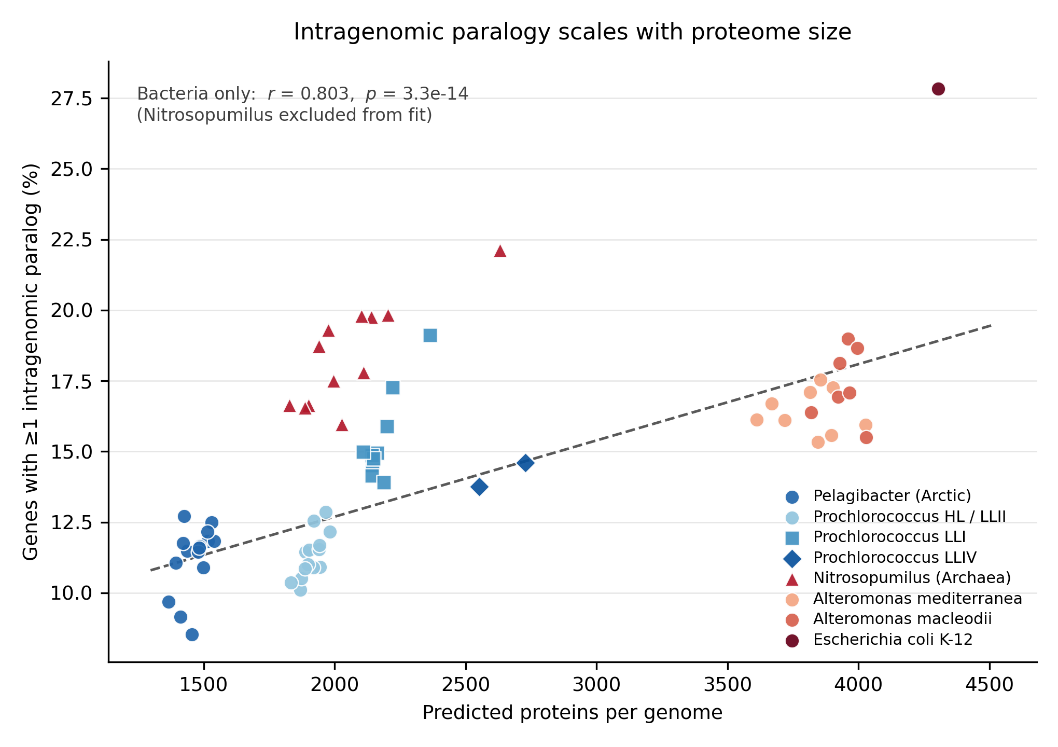

**Supplementary Information Figure 1. Intragenomic paralogy scales with proteome size.**  Proportion of genes with at least one paralog elsewhere in the same genome plotted against the number of predicted proteins. Each point represents one closed genome (n = 70; *Pelagibacter* (n = 15), *Prochlorococcus marinus* (n = 26), *Nitrosopumilus* sp. cultured isolates (n = 12, triangles), *Alteromonas mediterranea* (n = 9), *A. macleodii* (n = 7), and *E. coli* K-12 MG1655 (n = 1)). Paralogs were defined as protein pairs sharing ≥30% amino acid identity over an alignment spanning ≥70% of the length of both sequences, with self-hits and mobile elements excluded. The dashed line is the least-squares fit to the 58 bacterial genomes (r = 0.809, r² = 0.655, p = 1.5 × 10⁻¹⁴; slope 2.75 percentage points per 1,000 proteins); the archaeal lineage *Nitrosopumilus* was excluded from the fit and is shown for comparison.

| **Supplementary Information Table 1. Genomes analysed, assembly quality and intragenomic paralog content.** | | | | |  | |  | |  | |  | |  | |
| --- | --- | --- | --- | --- | --- | --- | --- | --- | --- | --- | --- | --- | --- | --- |
| CheckM2 completeness and contamination. Paralogy: BLASTP, E ≤ 1e-5, ≥30% (or ≥40%) amino acid identity over ≥70% of the length of both sequences, mobile elements excluded. Prokka annotation **throughout**. | | | | | | | | | | | | | | |
| **Lineage** | **Genome / assembly** | **Genome size (bp)** | **Contigs** | **CheckM2 completeness (%)** | | **CheckM2 contamination (%)** | | **Predicted proteins** | | **Mobile elements** | | **Paralogs ≥30% id (%)** | | **Paralogs ≥40% id (%)** |
| Pelagibacter (Arctic) | Uisw_090 | 1381496 | 1 | 98.54 | | 1.47 | | 1516 | | 1 | | 11.75 | | 4.55 |
| Pelagibacter (Arctic) | Uisw_092 | 1306717 | 1 | 98.28 | | 0.03 | | 1438 | | 0 | | 11.34 | | 3.06 |
| Pelagibacter (Arctic) | Uisw_094 | 1369821 | 1 | 96.25 | | 0.08 | | 1486 | | 0 | | 11.51 | | 3.57 |
| Pelagibacter (Arctic) | Uisw_099_02 | 1288341 | 1 | 99.87 | | 0.04 | | 1366 | | 0 | | 9.44 | | 2.27 |
| Pelagibacter (Arctic) | Uisw_101 | 1448726 | 1 | 99.8 | | 0 | | 1531 | | 2 | | 12.3 | | 5.23 |
| Pelagibacter (Arctic) | Uisw_104 | 1441579 | 1 | 96.9 | | 0.19 | | 1541 | | 0 | | 11.75 | | 4.87 |
| Pelagibacter (Arctic) | Uisw_106 | 1376435 | 1 | 97.8 | | 0.19 | | 1498 | | 0 | | 10.55 | | 3.6 |
| Pelagibacter (Arctic) | Uisw_113 | 1355470 | 1 | 99.47 | | 0.05 | | 1425 | | 0 | | 12.56 | | 3.86 |
| Pelagibacter (Arctic) | Uisw_116 | 1301732 | 1 | 99.77 | | 0.14 | | 1412 | | 0 | | 8.85 | | 2.12 |
| Pelagibacter (Arctic) | Uisw_121 | 1352740 | 1 | 98.92 | | 1.15 | | 1456 | | 0 | | 8.52 | | 1.79 |
| Pelagibacter (Arctic) | Uisw_127 | 1332340 | 1 | 99.56 | | 0.11 | | 1422 | | 0 | | 11.39 | | 3.02 |
| Pelagibacter (Arctic) | Uisw_130 | 1423135 | 1 | 99.95 | | 0 | | 1515 | | 2 | | 11.7 | | 3.64 |
| Pelagibacter (Arctic) | Uisw_134_02 | 1301899 | 1 | 99.86 | | 0.01 | | 1395 | | 0 | | 10.75 | | 2.87 |
| Pelagibacter (Arctic) | Uisw_136 | 1394121 | 1 | 99.81 | | 0.1 | | 1479 | | 0 | | 11.22 | | 3.04 |
| Pelagibacter (Arctic) | Uisw_137 | 1403101 | 1 | 99.85 | | 0.15 | | 1484 | | 1 | | 11.4 | | 3.3 |
| Prochlorococcus | AS9601__GCA_000015645.1 | 1669886 | 1 | 99.99 | | 0.08 | | 1889 | | 1 | | 11.44 | | 5.83 |
| Prochlorococcus | LG__GCA_047302905.1 | 1751940 | 1 | 99.99 | | 0.01 | | 1902 | | 0 | | 11.51 | | 6.36 |
| Prochlorococcus | MIT_0912__GCA_027359595.1 | 1898947 | 1 | 99.97 | | 0.01 | | 2145 | | 1 | | 14.51 | | 8.72 |
| Prochlorococcus | MIT_0913__GCA_027359525.1 | 1877877 | 1 | 99.99 | | 0.1 | | 2142 | | 0 | | 14.15 | | 7.84 |
| Prochlorococcus | MIT_0915__GCA_027359475.1 | 1988105 | 1 | 99.97 | | 0 | | 2187 | | 0 | | 13.9 | | 8.96 |
| Prochlorococcus | MIT_0917__GCA_027359575.1 | 1924674 | 1 | 100 | | 0.14 | | 2162 | | 1 | | 14.95 | | 9.35 |
| Prochlorococcus | MIT_0918__GCA_027359415.1 | 1789313 | 1 | 99.99 | | 0.06 | | 1944 | | 1 | | 10.91 | | 5.15 |
| Prochlorococcus | MIT_0919__GCA_027359375.1 | 1716770 | 1 | 100 | | 0.02 | | 1870 | | 0 | | 10.11 | | 4.39 |
| Prochlorococcus | MIT_1012__GCA_047302745.1 | 1897577 | 1 | 99.99 | | 0.25 | | 2221 | | 2 | | 17.26 | | 11.22 |
| Prochlorococcus | MIT_1013__GCA_027359395.1 | 2046712 | 1 | 100 | | 0.34 | | 2365 | | 0 | | 19.11 | | 12.9 |
| Prochlorococcus | MIT_1214__GCA_027359355.1 | 1934849 | 1 | 100 | | 0.06 | | 2143 | | 2 | | 14.85 | | 9.67 |
| Prochlorococcus | MIT_9107__GCA_047302795.1 | 1699808 | 1 | 100 | | 0.1 | | 1940 | | 0 | | 11.55 | | 6.39 |
| Prochlorococcus | MIT_9201__GCA_047305125.1 | 1685880 | 1 | 100 | | 0.06 | | 1942 | | 0 | | 11.69 | | 6.18 |
| Prochlorococcus | MIT_9215__GCA_000018065.1 | 1738790 | 1 | 100 | | 0.17 | | 1982 | | 1 | | 12.17 | | 6.61 |
| Prochlorococcus | MIT_9301__GCA_000015965.1 | 1641879 | 1 | 100 | | 0.02 | | 1873 | | 0 | | 10.52 | | 4.86 |
| Prochlorococcus | MIT_9302__GCA_047947085.1 | 1743611 | 1 | 100 | | 0.12 | | 1967 | | 1 | | 12.87 | | 6.92 |
| Prochlorococcus | MIT_9303__GCA_047302995.1 | 2686886 | 1 | 99.99 | | 0.2 | | 2728 | | 4 | | 14.61 | | 8.7 |
| Prochlorococcus | MIT_9312__GCA_000012645.1 | 1709204 | 1 | 100 | | 0.02 | | 1919 | | 1 | | 10.9 | | 5.68 |
| Prochlorococcus | MIT_9313__GCA_000011485.1 | 2410873 | 1 | 99.99 | | 0.02 | | 2552 | | 2 | | 13.76 | | 8.16 |
| Prochlorococcus | MIT_9515__GCA_000015665.1 | 1704176 | 1 | 100 | | 0.09 | | 1898 | | 0 | | 11.01 | | 5.58 |
| Prochlorococcus | NATL1A__GCA_000015685.1 | 1864731 | 1 | 99.99 | | 0.06 | | 2149 | | 1 | | 14.76 | | 9.31 |
| Prochlorococcus | NATL2A__GCA_000012465.1 | 1842899 | 1 | 99.99 | | 0.09 | | 2108 | | 0 | | 14.99 | | 9.68 |
| Prochlorococcus | PAC1__GCA_047304675.1 | 1927571 | 1 | 99.99 | | 0 | | 2199 | | 2 | | 15.89 | | 10.01 |
| Prochlorococcus | SB__GCA_047305225.1 | 1674013 | 1 | 100 | | 0.02 | | 1887 | | 0 | | 10.86 | | 5.99 |
| Prochlorococcus | SS51__GCA_047302855.1 | 1627281 | 1 | 100 | | 0.07 | | 1833 | | 1 | | 10.37 | | 5.29 |
| Prochlorococcus | pastoris_CCMP1986__GCA_047497125.1 | 1659596 | 1 | 99.98 | | 0.06 | | 1921 | | 0 | | 12.55 | | 6.56 |
| Nitrosopumilus (isolates) | Nitrosopumilus_adriaticus_NF5__GCF_000956175.1 | 1803090 | 1 | 99.89 | | 0.15 | | 2204 | | 5 | | 19.83 | | 12.14 |
| Nitrosopumilus (isolates) | Nitrosopumilus_cobalaminigenes_HCA1__GCF_013407145.1 | 1563356 | 1 | 99.11 | | 0.04 | | 1901 | | 2 | | 16.64 | | 8.95 |
| Nitrosopumilus (isolates) | Nitrosopumilus_koreensis_AR1__GCF_000299365.1 | 1639964 | 1 | 99.47 | | 0.54 | | 2027 | | 3 | | 15.96 | | 8.15 |
| Nitrosopumilus (isolates) | Nitrosopumilus_maritimus_SCM1__GCF_000018465.1 | 1645259 | 1 | 99.64 | | 0.04 | | 1941 | | 3 | | 18.73 | | 10.27 |
| Nitrosopumilus (isolates) | Nitrosopumilus_oxyclinae_HCE1__GCF_013407165.1 | 1585110 | 1 | 99.73 | | 0.1 | | 1886 | | 1 | | 16.55 | | 9.02 |
| Nitrosopumilus (isolates) | Nitrosopumilus_piranensis_D3C__GCF_000875775.1 | 1713078 | 1 | 99.81 | | 0.12 | | 2140 | | 5 | | 19.77 | | 11.52 |
| Nitrosopumilus (isolates) | Nitrosopumilus_sediminis_AR2__GCF_000299395.1 | 1690905 | 1 | 99.51 | | 0.13 | | 2110 | | 3 | | 17.8 | | 10.92 |
| Nitrosopumilus (isolates) | Nitrosopumilus_sp_K4__GCF_018128925.1 | 1645909 | 1 | 99.93 | | 0.02 | | 1977 | | 3 | | 19.3 | | 10.74 |
| Nitrosopumilus (isolates) | Nitrosopumilus_sp_S4__GCF_054515835.1 | 1701445 | 1 | 99.97 | | 0.19 | | 1996 | | 3 | | 17.51 | | 9.28 |
| Nitrosopumilus (isolates) | Nitrosopumilus_sp_SW__GCF_006740685.1 | 1499620 | 1 | 99.61 | | 0.08 | | 1827 | | 0 | | 16.64 | | 8.32 |
| Nitrosopumilus (isolates) | Nitrosopumilus_ureiphilus_PS0__GCF_013407185.1 | 2171418 | 1 | 99.76 | | 0.87 | | 2631 | | 6 | | 22.13 | | 14.67 |
| Nitrosopumilus (isolates) | Nitrosopumilus_zosterae_NM25__GCF_025998175.1 | 1775052 | 1 | 99.95 | | 0.12 | | 2102 | | 1 | | 19.8 | | 11.23 |
| Alteromonas mediterranea | MED_GCA_000020585.3 | 4480937 | 1 | 100 | | 0.7 | | 4028 | | 71 | | 15.95 | | 6.55 |
| Alteromonas mediterranea | MED_GCA_000439495.1 | 4397537 | 1 | 100 | | 0.92 | | 3855 | | 41 | | 17.54 | | 8.39 |
| Alteromonas mediterranea | MED_GCA_000439535.1 | 4442936 | 1 | 100 | | 0.21 | | 3897 | | 53 | | 15.58 | | 6.14 |
| Alteromonas mediterranea | MED_GCA_000439555.1 | 4395035 | 1 | 100 | | 0.13 | | 3844 | | 35 | | 15.33 | | 5.64 |
| Alteromonas mediterranea | MED_GCA_000439595.1 | 4438767 | 1 | 100 | | 1.56 | | 3902 | | 49 | | 17.26 | | 8.49 |
| Alteromonas mediterranea | MED_GCA_001886455.1 | 4344439 | 1 | 100 | | 0.27 | | 3668 | | 13 | | 16.69 | | 6.24 |
| Alteromonas mediterranea | MED_GCA_006517095.1 | 4343683 | 1 | 100 | | 0 | | 3719 | | 14 | | 16.11 | | 6.15 |
| Alteromonas mediterranea | MED_GCA_006517115.1 | 4294985 | 1 | 100 | | 0.39 | | 3611 | | 13 | | 16.12 | | 6.06 |
| Alteromonas mediterranea | MED_GCA_009759785.1 | 4512541 | 1 | 100 | | 0.01 | | 3815 | | 18 | | 17.09 | | 6.87 |
| Alteromonas macleodii | MAC_GCA_000172635.2 | 4653851 | 1 | 100 | | 1.06 | | 3960 | | 14 | | 18.98 | | 8.95 |
| Alteromonas macleodii | MAC_GCA_000299955.1 | 4601785 | 1 | 100 | | 0.05 | | 3923 | | 21 | | 16.94 | | 6.41 |
| Alteromonas macleodii | MAC_GCA_000299995.1 | 4480507 | 1 | 100 | | 0.12 | | 3819 | | 35 | | 16.38 | | 6.82 |
| Alteromonas macleodii | MAC_GCA_001562235.1 | 4575623 | 1 | 100 | | 0.12 | | 4029 | | 51 | | 15.51 | | 6.49 |
| Alteromonas macleodii | MAC_GCA_903772925.1 | 4665060 | 1 | 100 | | 0.05 | | 3966 | | 39 | | 17.09 | | 6.21 |
| Alteromonas macleodii | MAC_GCA_963691725.1 | 4670665 | 1 | 100 | | 0.24 | | 3995 | | 82 | | 18.66 | | 8.36 |
| Alteromonas macleodii | MAC_GCA_965277195.1 | 4671896 | 1 | 100 | | 0.03 | | 3929 | | 11 | | 18.12 | | 6.94 |
| Escherichia coli K-12 | MG1655 (U00096.3) | 4641652 | 1 |  | |  | | 4305 | |  | | 27.8 | |  |
